# Discovery of functional NLRs using expression level, high-throughput transformation, and large-scale phenotyping

**DOI:** 10.1101/2024.06.25.599845

**Authors:** Helen J. Brabham, Inmaculada Hernández-Pinzón, Chizu Yanagihara, Noriko Ishikawa, Toshiyuki Komori, Oadi N. Matny, Amelia Hubbard, Kamil Witek, Alexis Feist, Hironobu Numazawa, Phon Green, Antonín Dreiseitl, Naoki Takemori, Toshihiko Komari, Roger P. Freedman, Brian Steffenson, H. Peter van Esse, Matthew J. Moscou

## Abstract

Protecting crops from pests and diseases is vital for the sustainable agricultural systems needed for food security. Introducing functional resistance genes to enhance the plant immune system is an effective method of disease control, but identifying new immune receptors is time-consuming and resource intensive. We observed that functional immune receptors of the NLR class show a signature of high expression in uninfected plants across both monocot and dicot species. Here we show that this signature, combined with high throughput crop transformation, can be used to rapidly identify candidate NLRs from diverse plant species and validate pathogen resistance directly in crop plants. As a proof of concept, we generated a wheat transgenic library carrying 995 NLRs from 18 grass species. Screening the collection with the stem rust pathogen *Puccinia graminis*, which is a major threat to wheat production, we confirm 19 new resistance genes. This pipeline facilitates resistance gene discovery, unlocking a large gene pool of diverse and non-domesticated plant species and providing *in-planta* gene validation of disease resistance directly in crops.

## Introduction

Food insecurity is an increasing global crisis, as millions of people worldwide still face hunger and experience malnutrition (FAO, 2022; IPCC, 2022). Protecting plant health is vital for building the sustainable food systems needed to end hunger and poverty as plant diseases and pests cause major losses to crop yields worldwide (Anderson *et al*., 2004; Fisher *et al*., 2012; Savary *et al*., 2019). Plant pathogens have had an increasing impact in recent decades due to spread through international trade and travel, expansion of plant host ranges, and incursions of new pathogenic species and virulent strains. In addition, climate changes can alter the distribution of pests and diseases, contributing to new epidemics that can appear suddenly and spread rapidly (Bebber, 2015; Fisher *et al*., 2012; Hovmøller *et al*., 2022; Kettles & Luna, 2019; Shaw & Osborne, 2011). Thus, there is an urgent need for innovation and acceleration of novel methods to combat plant diseases.

One of the most effective and environmentally friendly methods of crop protection is to leverage the plant immune system for disease resistance to activate inherent defence responses against pathogens. Plants contain immune receptors that recognise pathogen invasion both outside and inside the cell. A major class of intracellular plant disease resistance genes encode nucleotide-binding domain leucine-rich repeat (NLR) proteins (Dodds & Rathjen, 2010). Breeding for disease resistance through transferring NLRs within and between plant species has proven successful against various pathogens. However, pathogens are constantly evolving and can overcome and evade NLRs, especially when NLRs are deployed over large areas in monocultures as individual genes. Introducing multiple NLRs into a single region of the genome through generating gene stacks can provide strong defence and increase the durability of these resistance genes (Hafeez *et al*., 2021; Luo *et al*., 2021; Van Esse *et al*., 2020; Wulff & Krattinger, 2022). Yet for this strategy to be effective, new and larger numbers of NLRs need to be identified as a limited number are available in modern cultivars. Wild relatives of crop species are often resistant to major agricultural pathogens, in a phenomenon described as non-host resistance (Schulze-Lefert & Panstruga, 2011). This resistance has been shown to involve NLRs particularly for closely related plant species (Panstruga & Moscou, 2020), but accessing the genes can be difficult using molecular genetic techniques. Large-scale projects to characterise NLRs from these genetically diverse sources are required to find new resistances against a wide range of pathogens (Bevan *et al*., 2017; Hafeez *et al*., 2021; Mascher *et al*., 2019; Wulff & Krattinger, 2022).

Since the first cloned resistance genes in the 1990s (Kourelis & Van Der Hoorn, 2018; Whitham *et al*., 1994), NLRs have been extensively studied via evolutionary and sequence-based analyses (Bailey *et al*., 2018; Kuang *et al*., 2004; Michelmore & Meyers, 1998; Shimizu *et al*., 2022; Van de Weyer *et al*., 2019; Yang *et al*., 2013). Furthermore, there is now knowledge of NLR structure (Förderer *et al*., 2022; Wang, Hu, *et al*., 2019; Wang, Wang, *et al*., 2019; Y. Yang *et al*., 2024), highly variable residues (Prigozhin & Krasileva, 2021; Sutherland *et al*., 2024), and sequence motifs important for function (Adachi *et al*., 2019). NLRs recognise pathogen infection by directly interacting with pathogen molecules or via recognising pathogen-induced modifications to plant host proteins (Cesari, Bernoux, *et al*., 2014; Van Der Biezen & Jones, 1998). Successful recognition results in defence responses to prevent the spread of infection, often resulting in localised cell-death (Saur *et al*., 2021). A few investigations have demonstrated that the presence or altered regulation of some NLRs cause deleterious effects: the presence of *Arabidopsis thaliana RPM1* has been shown to reduce silique and seed production (Tian *et al*., 2003), overexpression of *RPW8* can cause spontaneous cell death (Xiao *et al*., 2003), overexpression of *LAZ5* can cause lethality (Palma *et al*., 2010), and the lack of *PigmR* suppression in *Oryza sativa* causes a decrease in grain weight in rice (Deng *et al*., 2017). These observations, combined with the capability of NLRs to cause cell death, resulted in the pervasive idea that NLRs require strict regulation to control defence responses and are therefore transcriptionally repressed in uninfected plants (Balint-Kurti, 2019; Bergelson & Purrington, 1996; Brown, 2002; Brown & Rant, 2013; Howles *et al*., 2005; Karasov *et al*., 2017; Lai & Eulgem, 2018; Richard *et al*., 2018; Stokes *et al*., 2002; Tan *et al*., 2007).

Here, in our investigation of the NLR *Mla7* in barley, we found that multiple copies of *Mla7* are required for full complementation of host resistance to the barley powdery mildew pathogen *Blumeria hordei* (*Bh*) as well as non-host resistance to the wheat stripe rust pathogen *Puccinia graminis* f. sp. *tritici* (*Pgt*)and single copies of *Mla7* are insufficient for resistance. Previous work on *Mla3* also showed the requirement of multiple gene copies for resistance (Brabham *et al*., 2024), challenging the prevailing view that NLR expression must be maintained at a low level. Other characterised *Mla* alleles have been shown to directly recognise effectors from *Bh* (Saur *et al*., 2019), therefore we explored whether higher expression was correlated with functionality in NLRs. To investigate NLR expression, we assessed the expression levels of known characterised NLRs across six plant species of both monocots and dicots. In contrast to the literature, we observed an unexpectedly large number of NLRs are expressed in uninfected plants of these species. Furthermore, we examined known functional NLRs and found them to be present among highly expressed NLR transcripts at a high frequency.

We sought to exploit this observation to predict functional NLR candidates at scale and use high-throughput transformation and screening to greatly accelerate resistance gene discovery. As a proof of concept, we applied this approach to find novel resistance to stem rust of wheat, using high-efficiency wheat transformation (Ishida *et al*., 2015) to test NLRs directly in wheat. Stem rust is caused by the fungal pathogen *Pgt* and to date, 14 NLRs with efficacy against *Pgt* have been cloned (Arora *et al*., 2019; Chen *et al*., 2018; Luo *et al*., 2022; Mago *et al*., 2015; Periyannan *et al*., 2013; Saintenac *et al*., 2013; Steuernagel *et al*., 2016; Zhang *et al*., 2021; Zhang *et al*., 2023; Zhang *et al*., 2017). This limited set of *Pgt* resistance genes has proved insufficient to control disease outbreaks and for the development of wheat varieties with durable resistance (Hafeez *et al*., 2021; Wulff & Dhugga, 2018; Wulff & Krattinger, 2022). We generated a transgenic array of 995 NLRs from 18 diverse grass species and identified 19 new NLRs that conferred resistance to stem rust in wheat. This wheat transgenic array is a valuable resource that can be repeatedly screened with new pathogens and pathogen races to identify additional effective NLRs in the collection. The functional NLR expression signature found across plant species in our study implies that this approach could be replicated in other crops and their wild relatives, to identify new resistance genes. Generating a pan-plant NLR resource—combining functional resistance genes and informatics tools—would empower the discovery of resistance genes against a wide variety of pathogens and production of disease-resistant crops.

## Results

### The NLR *Mla7* requires multiple copies to confer resistance to barley powdery mildew and wheat stripe rust

In barley, alleles of the NLR *Mla* are known to confer resistance to barley powdery mildew caused by *Bh*. Through transgene complementation in a susceptible wild-type background, we confirmed that *Mla7* confers resistance to *Bh* isolate CC148 carrying the recognised effector *AVR_a7_* (**Fig. 1a**). We observed that single insertions of the transgene were insufficient to complement the resistance phenotype. Two independent single copy insertion lines of *Mla7* (T1-1 and T1-3) were susceptible to *Bh* isolate CC148 (*AVR_a7_*), whereas two multicopy insertion lines (T1-4 and T1-8) expressed resistance (**Fig. 1a**). To confirm race-specificity of resistance, we inoculated the transgenic lines carrying *Mla7* with 13 diverse *Bh* isolates that vary for presence of the recognised effector *AVR_a7_*. Resistance to *Bh* isolates with *AVR_a7_* was only observed in *Mla7* multicopy insert lines, although considerable variation was observed within and between transgenic families (**Fig. 1c**).

**Figure 1.**
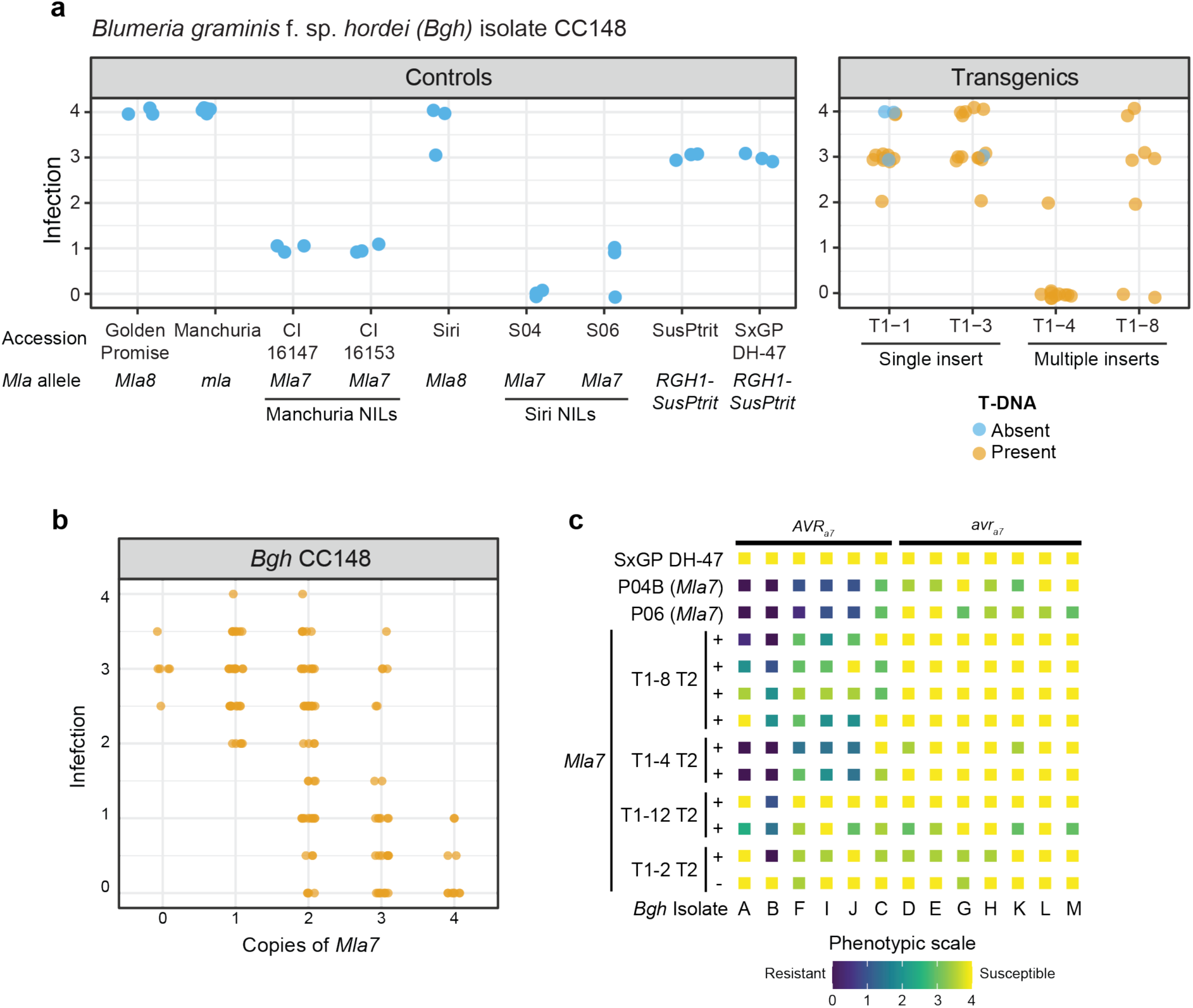
Multiple copies of *Mla7* are required to confer resistance to barley powdery mildew caused by *Blumeria hordei* (*Bh*). Barley powdery mildew susceptible barley cv. SxGP DH-47 was transformed with *Mla7* driven by the *Mla6* promoter-5’UTR and *Mla6* 3’UTR-terminator. Three single copy insert lines (T1-1, T1-2, and T1-3) and three multiple copy insert lines (T1-4, T1-8, and T1-12) were identified for *Mla7*. Resistance to *Bh* isolates carrying *AVR_a7_* was observed in transgenic barley lines carrying multiple copies of *Mla7.* Specific recognition of the effector *AVR_a7_* was retained across transgenic lines. **A**) Infection phenotypes for presence (+) or absence (-) of T-DNA for *Mla7* T_1_ families and controls inoculated with *Bh* isolate CC148. Presence or absence of T-DNA is shown in orange and blue, respectively. All phenotypes are on a scale from 0.0 to 4.0 in 0.5. Transparency and jittering were used to visualize multiple overlapping data points. **B**) Multiple copies, not single copies, of *Mla7* under its native promoter/terminator are required to confer full resistance to *Bh* isolate CC148. Individual phenotypes from an F_2_ population with varying copy number of *Mla7* transgene copies derived from a cross of two single insertion transgenic lines (T1-117 and T1-121). **C**) Multicopy lines carrying *Mla7* driven by the *Mla6* promoter/terminator confers race-specific resistance to barley powdery mildew (*Bh*). Controls include SxGP DH-47 and near-isogenic lines in the Pallas (*Mla8*) genetic background: P04B (*Mla7*), and P06 (*Mla7*). *Bh* isolates are ordered based on the presence (*AVR_a7_*) or absence (*avr_a7_*) of the effector recognised by *Mla7.* Isolates include 3-33 (A), Race I (B), X-4 (C), I-167 (D), K-200 (E), M-236 (F), Z-6 (G), C-132 (H), 120 (I), R86/1 (J), K-3 (K), KM18 (L), and MN-B (M). All experiments were performed twice with similar results, data shown is the average of these experiments.

*Mla* alleles are known to recognise multiple pathogens, with *Mla8* conferring resistance to *Bh* and wheat stripe rust caused by *Puccinia striiformis* f. sp. *tritici* (*Pst*) (Bettgenhaeuser *et al*., 2021) and *Mla3* conferring resistance to *Bh* and *Magnaporthe oryzae* (rice blast) (Brabham *et al*., 2024). In previous work, *Mla7* was shown to be in complete genetic coupling with *Rps7*, which confers resistance to *Pst*, and they are hypothesised to be the same gene (Bettgenhaeuser *et al*., 2021). Using the *Mla7* transgenic lines, we confirmed that *Mla7* also confers resistance to *Pst* (**Supplemental Fig. 1a**). In addition, like observed with *Bh*, multiple insertions of the *Mla7* transgene were required for *Pst* resistance (**Supplemental Fig. 1b**).

To determine whether the *Mla7* promoter was sufficient for expression of *Mla7*, we developed single copy transgenic lines expressing *Mla7* under its native promoter. Preliminary results found T_1_ families to be susceptible to *Bh*, therefore we reasoned that multiple copies may be required for function. We crossed two T_1_ families (T1-117 and T1-121) to develop an F_2_ population segregating for 0 to 4 copies of *Mla7*, to mitigate the challenges associated with multisite insertion and transgene silencing in multicopy lines. Higher order copies were required for resistance to *Bh*, as only transgenic lines carrying 2 or more copies showed resistance to *Bh* isolate CC148 (*AVR_a7_*) (**Fig. 1a**). Full recapitulation of native *Mla7*-mediated resistance was only observed in lines with four copies of *Mla7* (**Fig. 1b**). *Mla7* natively exists with three identical copies in the haploid genome of barley cv. CI 16147 (**Supplemental Fig. 2**), therefore the observation that multicopy insertions are required for resistance supports the hypothesis that a specific threshold of expression is required for function.

### Functional NLRs exhibit high steady-state expression levels

We next explored the expression levels of characterised functional NLRs from different plant species. High expression for functional NLRs was found in monocot species, using sequencing data from uninfected plant tissue. The barley (*Hordeum vulgare*) resistance genes *Rps7*/*Mla8* in the cv. Golden Promise and *Rps7*/*Mla7* in accessions CI 16147 and CI 16153 against wheat stripe rust (*Pst*) are present in highly expressed transcripts (**Fig. 2a**). The *Aegilops tauschii*-derived wheat stem rust (*Pgt*) resistance genes *Sr46*, *SrTA1662*, and *Sr45* are also present in highly expressed NLR transcripts across accessions (**Fig. 2b**). High expression of functional NLRs was also observed in dicot species. In *Arabidopsis thaliana*, highly expressed NLR transcripts are enriched with known and characterised resistance genes (**Fig. 2c; Supplemental Data 1**). The most highly expressed NLR in ecotype Col-0 is *ZAR1* which recognises bacterial pathogen effectors from *Pseudomonas syringae* and *Xanthomonas campestris* pv. *campestris* (**Fig. 2c; Supplemental Data 2**). Other highly expressed functional NLRs include the *RPP* alleles against *Hyaloperonospora arabidopsidis*, *WRR4* which provides resistance to *Albugo candida* and the defence response mediator *SNC1*. This observation is consistent across ecotypes, as the *RPP* and *WRR* alleles are also highly expressed in Ler-0, Sf-2, and Ws-0 seedlings. The most highly expressed NLR in Sf-2 and Ws-0 is an allele of *RLM3* which provides resistance to *Leptosphaeria maculans* and the necrotrophic pathogens *Botrytis cinerea*, *Alternaria brassicicola,* and *Alternaria brassicae* (**Supplemental Data 2)**. As a model species, *A. thaliana* contains a large complement of known and characterised functional NLRs. Using gene descriptions from Araport11, we observed a high enrichment of functional NLRs among highly expressed genes. Within the accession Col-0, we found that known NLRs are significantly enriched in the top 15% of expressed NLR transcripts compared to the lower 85% (*X^2^*(1, *N* = 616) = 4.2979, *p* = 0.038). Using a non-redundant set of the highest expressed transcript for each NLR, we found the top 14% of expressed NLR transcripts is enriched for known NLRs (*X^2^* (1, *N* = 141) = 4.5767, *p* = 0.032). NLRs in the top 15% are in NLR classes containing coiled-coil, nucleotide-binding-site, leucine-rich-repeat, and Toll/interleukin1 receptor domains without additional non-canonical domains (CNL, NL, TN, TNL, and TNLT; **Supplemental Fig. 3, Supplemental Fig. 4**). Overall, the expression level of the most highly expressed NLR is above the median and mean expression level for all genes in *A. thaliana* accession Col-0, confirming that NLRs are expressed in uninfected plants (**Supplemental Fig. 3, Supplemental Fig. 4**).

**Figure 2.**
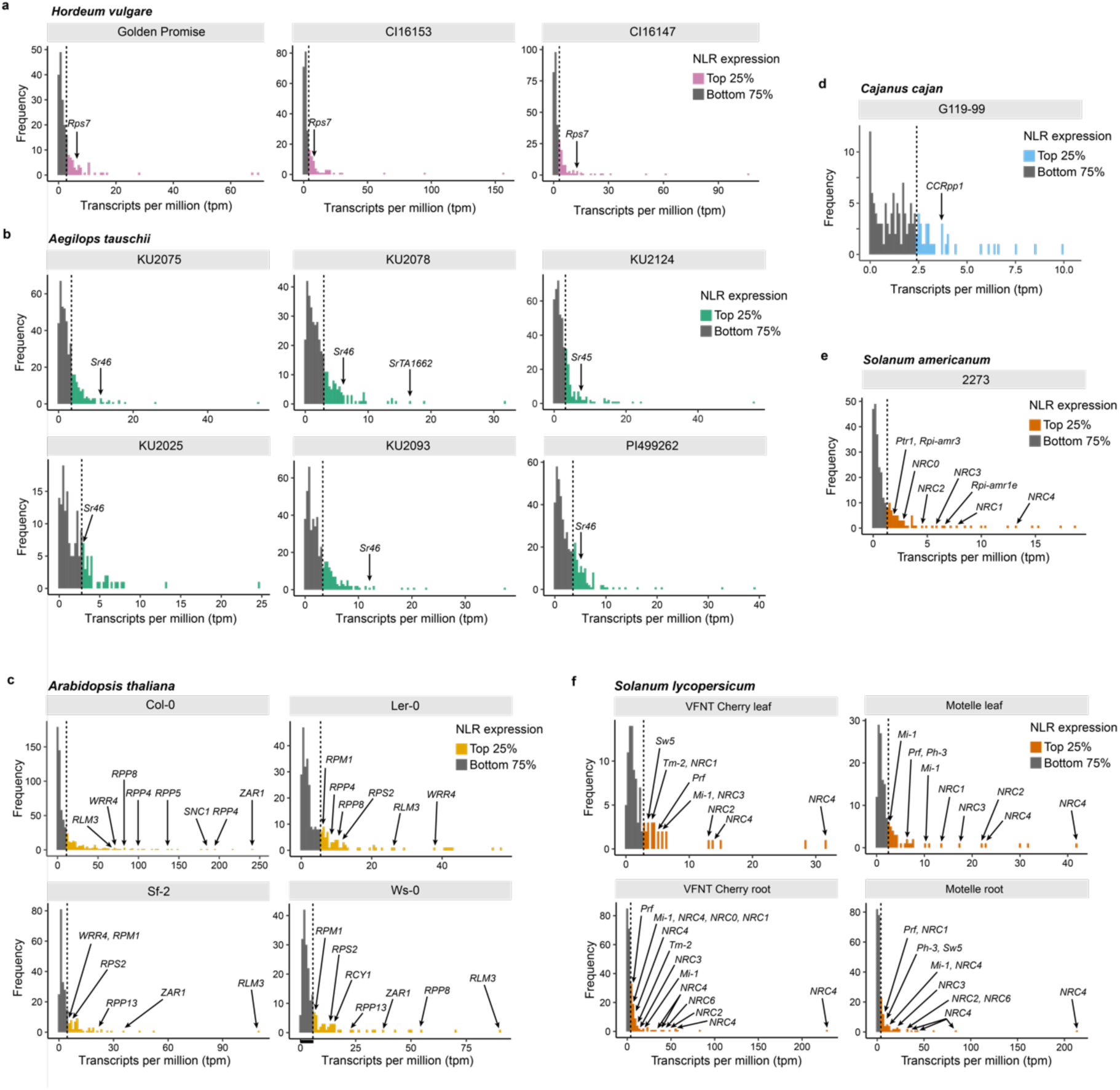
Functional NLRs are highly expressed in the *de novo* assembled transcriptome of unchallenged plants of *Aegilops*, *Cajanus*, *Arabidopsis*, *Solanum,* and *Hordeum* species. Known functional NLRs from each species are amongst the most highly expressed NLR transcripts from unchallenged plant tissue (**Supplemental Data 2**). Transcript abundance was estimated from self-aligned RNAseq data measured in transcripts per million (TPM). **A)** Transcript abundance of *NLRs* from the *de novo* assembled transcriptome of *Hordeum vulgare* accessions Golden Promise, CI16153, and CI16147. The expression of the functional resistance gene *Rps7* against *Blumeria graminis* and *Puccinia striiformis* f. sp. *tritici*. **B)** Transcript abundance of *NLRs* from the *de novo* assembled transcriptome of *Aegilops tauschii* accessions KU2025, KU2075, KU2078, KU2093, KU2124, and PI499262. The expression of the functional resistance genes *Sr45*, *Sr46*, and *SrTA1662* against *Puccinia graminis* f. sp. *tritici*. **C)** Transcript abundance of NLRs from *Arabidopsis thaliana* accession Col-0, Ler-0, Sf-2, and Ws-0. The expression of the functional resistance genes *RPP4*, *RPP5*, *RPP8, RPP13* to downy mildew (caused by *Hyaloperonospora arabidopsidis*); *WRR4* against white rust (*Albugo candida*); *ZAR1* against *Pseudomonas syringae* and *Xanthomonas campestris* pv. *campestris; RPS2* and *RPM1* against *Pseudomonas syringae*; *RCY1* against Cucumber mosaic virus (CMV); and alleles of *RLM3* against grey mould (*Botrytis cinerea*), dark leaf spot of cabbage (*Alternaria brassicicola*) and dark spot of crucifers (*Alternaria brassicae*). **D)** Transcript abundance of NLRs from *Cajanus cajan* accession G119-99. The expression level of *CCRpp1* which confers resistance to Asian soybean rust (*Phakopsora pachyrhizi*) (Kawashima *et al*., 2016). **E)** Transcript abundance of NLRs from the *de novo* assembled transcriptome of *Solanum americanum* accession SP2273. The expression of the functional resistance genes *Rpi-amr1* and *Rpi-amr3* to late blight (*Phytophthora infestans*). An allele of *Ptr1*, which recognises *Pseudomonas syringae* pv. *tomato* and *Ralstonia pseudosolanacearum* in *Solanum lycopersicoides,* is also shown. Helper NLRs annotated as *NRC1*, *NRC2*, *NRC3*, *NRC4*, and *NRC0* as defined by Kourelis *et al*., (2021). *NRC6* is not present in the dataset. **F)** Transcript abundance of NLRs from *Solanum lycopersicum* cultivars VFNT Cherry and Motelle carrying *Mi-1* with resistance to root-knot nematodes (*Meloidogyne* spp.), the potato aphid (*Macrosiphum euphorbiae*), and the sweet potato whitefly (*Bemisia tabaci*). *Mi-1* is highly expressed in both cultivars in both leaf and root tissue. The expression level of additional known functional NLRs for: *Tm-2* for resistance to tobamoviruses including *Tomato Mosaic Virus* (*ToMV*) and *Tobacco Mosaic Virus* (*TMV*); *Prf* for resistance to *Pseudomonas syringae* pv. *tomato*; *Sw5* for resistance to a broad range of viruses across paralogs; and *Ph-3* for resistance to *Phytophthora infestans*. Helper NLRs annotated as *NRC1*, *NRC2*, *NRC3*, *NRC4*, *NRC6*, and *NRC0* as defined by Kourelis *et al*., (2021).

In other plant species, known NLRs have been identified using methods including map-based cloning, association genetics, RenSeq (Arora *et al*., 2019; Jupe *et al*., 2013; Steuernagel *et al*., 2016), and other bioinformatic tools (Steuernagel *et al*., 2020). The NLRs *CcRpp1* from *Cajanus cajan* and *Rpi-amr1* from *Solanum americanum*, identified via these methods, were found to be present in highly expressed NLRs in the respective species (**Fig. 2d & 2e**). The tomato NLR *Mi-1*, previously confirmed by positional cloning, provides resistance to potato aphid and whitefly in foliar tissue and the root-knot nematode in the roots. We found that *Mi-1* is highly expressed in both the leaves and roots of the resistant cultivars Motelle and VFNT Cherry, alongside the additional characterised NLRs *Tm-2*, *Prf*, *Sw5*, and *Ph-3* (**Fig. 2f; Supplemental Data 2**).

*Rpi-amr1* and *Mi-1* are dependent on additional NLRs for function and are present in a wider network described in *Solanaceae* species (Witek *et al*., 2021; Wu *et al*., 2017). NLRs that recognise pathogen products or host modifications directly are described as ‘sensor’ NLRs and these partner with ‘helper’ NLRs that facilitate immune signalling (Adachi & Kamoun, 2022). Known ‘helper’ NLRs, designated with the prefix NRC (NLR required for cell death) in the *Solanaceae*, are also highly expressed NLRs (**Fig. 2e & f; Supplemental Data 2**). In addition, many ‘helper’ NLRs display tissue specificity; *NRC6* is highly expressed in the roots but not the leaves of tomato cvs. VFNT Cherry and Motelle, and *NRC0* is highly expressed in the roots of cv. VFNT Cherry but lowly expressed in the leaves of both cultivars, and in the roots of cv. Motelle (**Fig. 2f**). Therefore, these results show the importance of investigating the appropriate plant tissue based on the tissue specificity of the plant pathogen and indicates tissue specificity of resistance.

### The most highly expressed isoform of *Rpi-amr1* is the functional NLR

Multiple isoforms of each NLR are present in transcriptomes and while the function of alternative splicing and isoform variation across NLRs is broadly uncharacterised, alternatively spliced variants of a few NLRs have been shown to modulate defence. For example, presence of the full-length and truncated *A. thaliana RPS4* transcripts (Zhang & Gassmann, 2003), the tobacco (*Nicotiana glutinosa*) *N* gene (Dinesh-Kumar & Baker, 2000), and *Medicago truncatula RCT-1* (Tang *et al*., 2013) are required for resistance function. In addition, only one transcript variant of *Oryza sativa RGA5* confers resistance (Cesari *et al*., 2013) and variants of *Triticum aestivum TaYRG1* have been implicated in modulation of defence pathways (Zhang *et al*., 2022). To assess the role of different gene isoforms, we investigated *Rpi-amr1* from *S. americanum* which provides resistance to *Phytophthora infestans* (Witek *et al*., 2021). Different isoforms of *Rpi-amr1* are present in the assembled transcriptome of *S. americanum* accession SP2273 at varied expression levels. *Rpi-amr1* isoforms show sequence variation, including the presence/absence of the final exon (**Supplemental Fig. 5**). Expressing the different isoforms under the same NRC4 promoter in transient assays in *N. benthamiana*, we found that the most abundant isoform, *i3,* provides resistance to *P. infestans* (**Fig. 3**). The isoform *i3* contains all exons as compared to the published sequence. Isoform *i1* also confers resistance to *P. infestans* as it contains all exons and is 99.3% similar to *i3*. Other isoforms present at lower TPM levels in the transcriptome confer a reduced level of resistance to *P. infestans*. Transcript variants may be due to alternative splicing, the presence of paralogs, or transcript assembly. As the most highly expressed isoform of *Rpi-amr1* is the functional transcript, this indicates that selecting the highest expressed transcript is a feasible approach to select functional variants for other NLRs.

**Figure 3.**
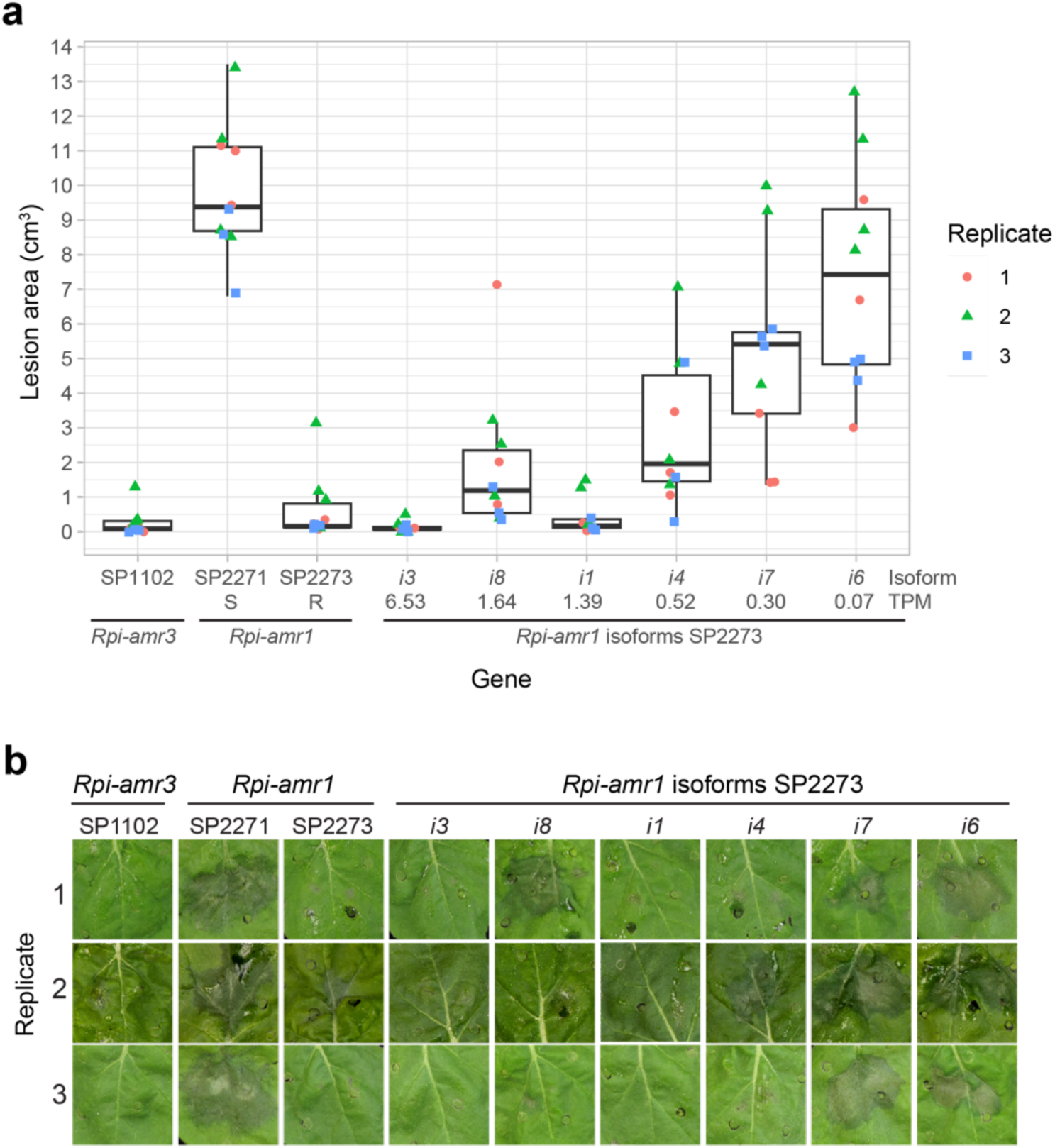
The most highly expressed isoform of *Rpi-amr1* provides resistance to *P. infestans*. Controls of resistant *Rpi-amr3* from accession SP1102, susceptible *Rpi-amr1* from accession SP2271, and resistant *Rpi-amr1* from accession SP2273 included. *Rpi-amr1* from accession SP2273 are present in descending order of expression level: *i3* at 6.53 TPM, *i8* at 1.64 TPM, *i1* at 1.39 TPM, *i4* at 0.523343 TPM, *i7* at 0.30 TPM, and *i6* at 0.07 TPM. **A)** Point plot and box plots of lesion area of *P. infestans* infection in transient assays in *N. benthamiana* using different isoforms of *Rpi-amr1*. Individual biological replicates in each replicate indicated with different coloured shapes (1: red circle, 2: green triangle, 3: blue square). **B)** Representative photographs of lesions across replicates.

### Building and testing of NLR arrays using the signature of high expression, high-throughput transformation, and large-scale phenotyping

We tested if we could use the signature of higher expression level in unchallenged tissue to mine functional NLRs from diverse plant germplasm. We sought to develop a pipeline to generate an array of candidate functional NLRs that could be tested using high-throughput screening. As a proof of concept, we hypothesised we could use this approach to identify new NLRs from diverse grass species for use against cereal rust pathogens which are a major threat to global wheat production. NLRs providing resistance to rust have been previously characterised from close relatives of wheat including *Aegliops* species (Arora *et al*., 2019; Periyannan *et al*., 2014; Steuernagel *et al*., 2016). Therefore, we investigated in total 30 accessions of *Aegilops bicornis, Aegilops longissima, Aegilops searsii,* and *Aegilops sharonensis* to capture the intraspecific and interspecific genetic diversity present in this genus (**Fig. 4a**; **Supplemental Table 1**). To represent the breadth of diversity across the grasses, we included several species of wheat relatives across the *Triticeae* and *Poeae* through to the distant relatives spanning the *Pooideae*—many of which have not previously been mined for rust resistance.

**Figure 4.**
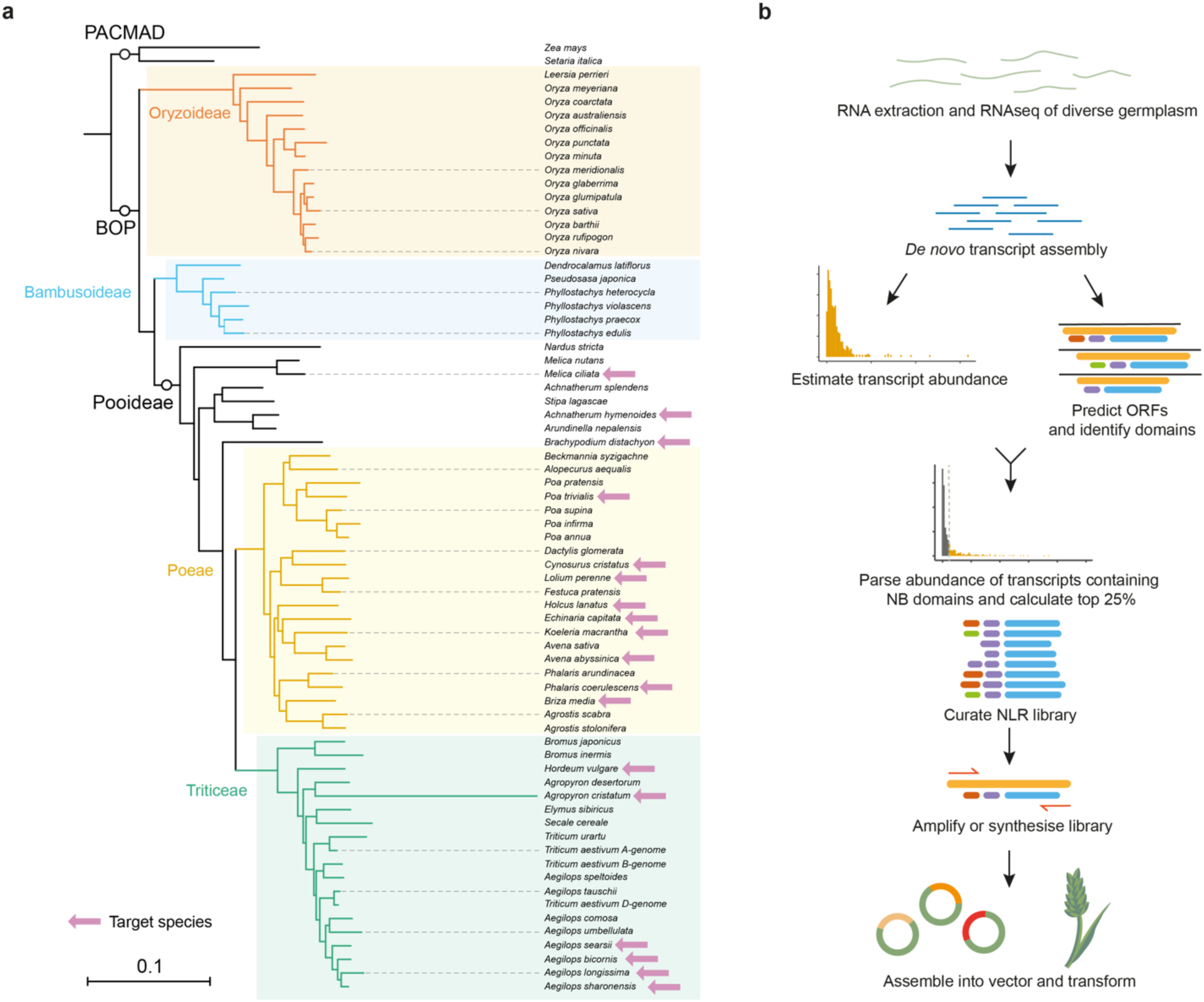
A pipeline for rapid identification of NLRs from diverse grass species. **A)** Phylogenetic tree of grass species within the PACMAD and BOP clades of the Poaceae. Species used for NLR discovery are indicated with pink arrows. **B)** Graphical representation of the pipeline. Identification of highly expressed NLRs from RNAseq data. The expression of transcripts containing an NB domain was used to identify the top 25% of expressed NLRs. Primers were designed on the open reading frame (ORF) and the coding sequence (CDS) were amplified via PCR and assembled into the transformation vector.

We identified candidate functional NLRs from a total of 69 accessions of 18 plant species (**Supplemental Table 1**). We excluded NLRs from the MIC1 clade, as this clade is enriched with NLRs with integrated domains that require an additional NLR to function together as a pair (Bailey *et al*., 2018; Cesari, Kanzaki, *et al*., 2014; Wang *et al*., 2013). The corresponding clade of paired ‘helper’ NLRs was retained as it is unknown if these NLRs can pair with multiple MIC1 NLRs or function in interspecific NLR pairings. In *A. thaliana,* the top 15% of NLR transcripts is enriched with known NLRs and we expanded this threshold in our pipeline to the top 25% to ensure broad capture of functional NLRs. The 25% threshold captures 61% of known NLRs in *A. thaliana* excluding NLRs with additional integrated domains. This threshold also encompasses known NLRs across plant species, especially monocots (**Fig. 2**). We selected the most highly expressed isoform under the hypothesis this is the functional variant. Using high-efficiency, high-throughput *Agrobacterium*-mediated transformation of the wheat cultivar Fielder (Ishida *et al*., 2015), a total of 6,260 transformation events generated 5,177 independent T_1_ families for 995 NLR constructs to create an array of transgenic wheat (**Fig. 4b; Supplemental Data 3**). This array also includes the controls of known NLRs conferring stem rust resistance: *Sr33* (Periyannan *et al*., 2013), *Sr35* (Saintenac *et al*., 2013), and *Sr50* (Mago *et al*., 2015).

### A total of 19 new NLRs confer resistance to wheat stem rust

We interrogated the transgenic wheat array for resistance to *Pgt* in greenhouse seedling assays using *Pgt* race QTHJC, a highly virulent isolate on wild-type Fielder (**Fig. 5a & 5b; Supplemental Data 4; Supplemental Table 2**). A total of 19 NLRs provided resistance to *Pgt* race QTHJC when expressed in Fielder. Resistance was observed in two or more independent T_1_ families for six NLRs and in one independent T_1_ family for 13 of the NLRs. Control lines carrying the known resistance gene *Sr50* were resistant to *Pgt* race QTHJC (**Fig. 5a; Supplemental Data 4; Supplemental Table 2**). All other remaining NLRs showed a susceptible phenotype (**Fig. 5a**; **Supplemental Fig. 6**; **Supplemental Data 4**), including those carrying controls *Sr33* and *Sr35*. The susceptible phenotype observed in *Sr33* transgenics may represent insufficient complementation by the transgene. Transgenic lines carrying *Sr35* were previously shown to be susceptible to *Pgt* race QTHJC, as this isolate is virulent on *Sr35* (Luo *et al*., 2021; Rouse *et al*., 2011; Saintenac *et al*., 2013).

**Figure 5.**
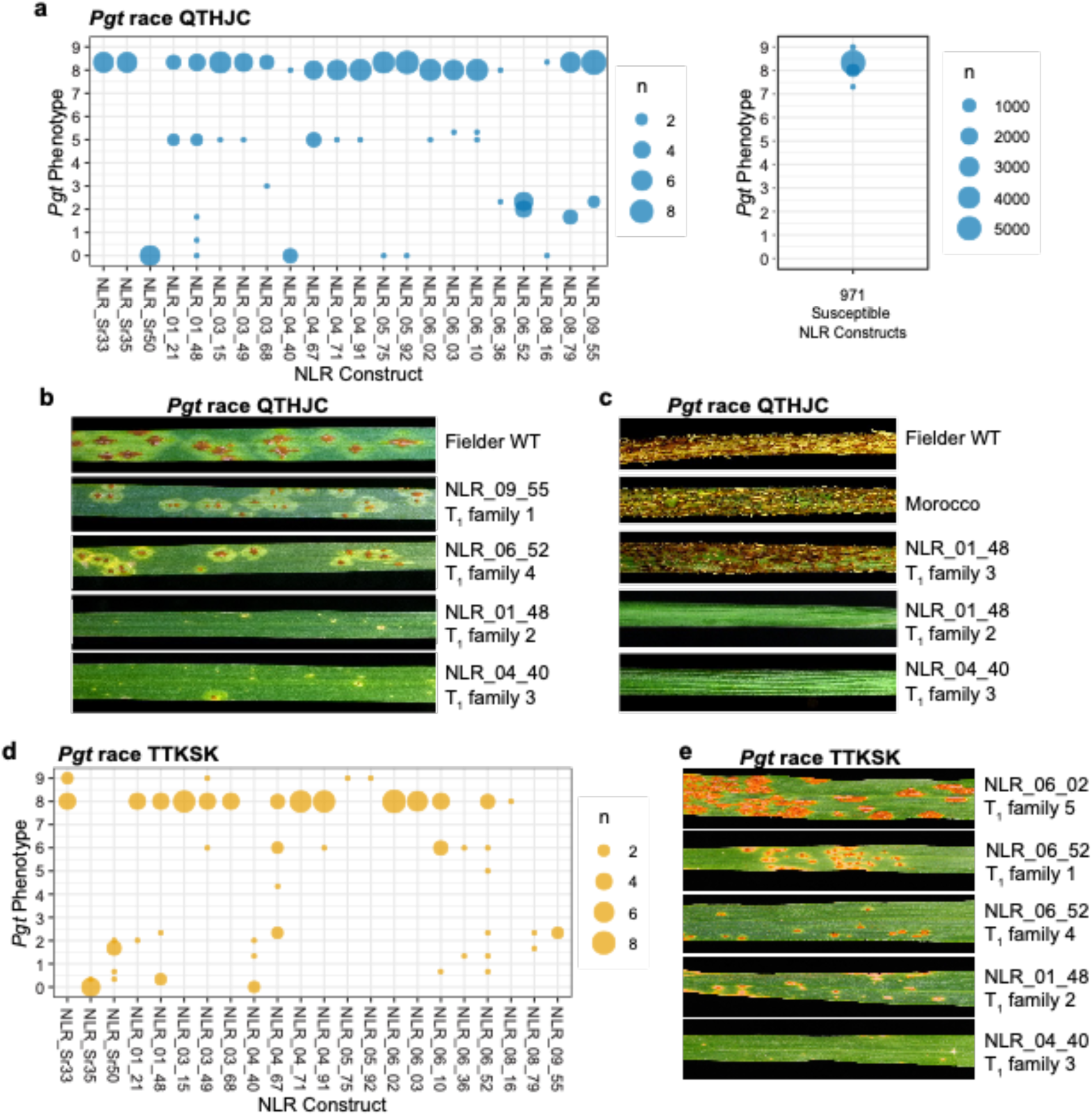
Identified NLRs provide resistance to *Pgt* races QTHJC and TTKSK. **A)** A total of 19 NLRs conferred resistance against *Pgt* race QTHJC (**Supplemental Data 4 and 5**). Phenotypic scores from individuals within T_1_ families from each construct inoculated with *Pgt* race QTHJC plotted on a weighted and transformed Stakman scale from completely resistant (0) to completely susceptible (9). Circle size indicates number of individuals having each phenotypic score. Individuals from the other 971 susceptible NLRs shown on the right-hand side were susceptible to *Pgt* and exhibited phenotypic scores similar to wild-type Fielder. **B)** Seedling leaves of T_2_ individuals from T_1_ families infected with *Pgt* race QTHJC under greenhouse conditions. From *top* to *bottom*: wild-type Fielder susceptible control, resistant individual from NLR_09_55 independent T_1_ family 1, resistant individual from NLR_06_52 independent T_1_ family 4, resistant individual from NLR_01_48 independent T_1_ family 2, and resistant individual from NLR_04_40 independent T_1_ family 3. **C)** Stem sections of individuals infected with *Pgt* race QTHJC under field conditions. From *top* to *bottom*: wild-type Fielder susceptible control, susceptible wheat control of cv. Morocco, susceptible individual from segregating family from NLR NLR_01_48 T_1_ family 3, resistant individual from NLR NLR_01_48 T_1_ family 2, and resistant individual from NLR NLR_04_40 T_1_ family 3. **D)** Nine NLRs conferred resistance against *Pgt* race TTKSK. Individuals with resistance against *Pgt* race QTHJC screened with *Pgt* race TTKSK. Phenotypes scored on a weighted Stakman scale from completely resistant (0) to completely susceptible (9). Circle size indicates number of individuals having each phenotypic score. Individuals from T_1_ families of NLRs NLR_01_21, NLR_01_48, NLR_04_40, NLR_04_67, NLR_06_10, NLR_06_36, NLR_06_52, NLR_08_79 and NLR_09_55 showed resistant phenotypes to *Pgt* race TTKSK. **E)** Seedling leaves of individuals from T_1_ families inoculated with *Pgt* race TTKSK under greenhouse conditions. From *top* to *bottom*: susceptible individual of NLR NLR_06_02 independent T_1_ family 5, resistant individual of NLR_06_52 independent T_1_ family 1, resistant individual of NLR_06_52 independent T_1_ family 4, resistant individual of NLR_01_48 independent T_1_ family 2, and resistant individual of NLR_04_40 independent T_1_ family 3.

Seed from individual resistant transgenic lines was bulked following the first round of inoculations and T_2_ families were tested to confirm efficacy in the field. A total of 17 of the 19 resistant NLRs were tested under field conditions, with 13 additional NLRs included as controls for comparison. T_2_ families of four NLRs showed resistance to *Pgt* race QTHJC in the field, (**Fig. 5c**; **Supplemental Fig. 7; Supplemental Data 5**). The reduced resistance compared to seedling assays observed from other NLR constructs may be due to developmental-dependent resistance responses.

The resistant transgenic lines conferring resistance to *Pgt* race QTHJC were inoculated with the widely virulent *Pgt* race TTKSK to assess the breadth of pathogen recognition. Nine of the 19 NLRs also conferred resistance to *Pgt* race TTKSK (**Fig. 5d and 5e; Supplemental Data 4; Supplemental Table 2**). Ten of the 19 NLRs displayed race-specific resistance to *Pgt* race QTHJC that was overcome by *Pgt* race TTKSK (**Fig. 5d**; **Supplemental Data 4; Supplemental Table 2**). To further test for pathogen recognition specificity and to exclude the possibility of constitutive activation of defence in transgenic lines, a sample of 5 NLRs conferring resistance to stem rust were tested additionally with *Puccinia triticina*, the fungal wheat leaf rust pathogen. The known leaf rust resistance gene, *Lr21* (Huang *et al*., 2009), was included as a resistant control (**Supplemental Fig. 8**). All transgenic lines carrying the selected 5 NLRs were susceptible to leaf rust, whereas *Lr21* showed resistance. Therefore, NLRs within the transgenic array retain pathogen recognition specificity and the resistance phenotypes observed are not due to constitutive defence activation by the transgenes.

### NLR hits are present across diverse phylogenetic clades

We investigated the distribution of the functional NLR hits against *Pgt* using a previously annotated phylogenetic tree of NLRs from grass species (Bailey *et al*., 2018). We found that the NLR hits are present across different phylogenetic clades (**Supplemental Table 2; Supplemental Fig. 9**). Six hits are present in the C17 clade which contains the known rust resistance genes *Lr10*, *Sr33*, and *Sr35* (Bailey *et al*., 2018). The C17 clade contains the NLRs *Sr35* and *Mla* which directly bind pathogen effectors (Förderer *et al*., 2022; Saur *et al*., 2019), and we hypothesise these *Pgt* hits also function as single NLRs via direct recognition. Two hits are present in the C24 clade which contains the known leaf rust resistance genes *Lr1* and *Lr21*. Five hits are derived from the C7 NLR clade enriched with known paired NLRs with a ‘helper’ function from the MIC1 clade (Bailey *et al*., 2018).

## Discussion

Our observation that a remarkable number of functional NLRs are highly expressed in uninfected plants provides a useful tool for facilitating large-scale resistance gene mining to combat major plant pathogens. In our proof-of-concept study, the cross-species transfer of single NLRs—including NLRs from species never genetically investigated for resistance to wheat pathogens—resulted in functional resistance in wheat. The NLR array generated from a wide range of grass species provided 19 new resistance genes with efficacy against *Pgt,* including nine NLRs with resistance to the widely virulent *Pgt* race TTKSK (Ug99 lineage). This method greatly accelerates NLR discovery for *Pgt* resistance, as only 14 NLRs with efficacy against *Pgt* have been cloned to date (Arora *et al*., 2019; Chen *et al*., 2018; Luo *et al*., 2022; Mago *et al*., 2015; Periyannan *et al*., 2013; Saintenac *et al*., 2013; Steuernagel *et al*., 2016; Zhang *et al*., 2021; Zhang *et al*., 2023; Zhang *et al*., 2017). Furthermore, the resulting NLR array is a valuable resource that can be repeatedly interrogated *in planta* against other important pests and diseases to find new effective resistance genes.

Further investigation is required as to elucidate the mechanism of recognition of these new *Pgt* resistant NLRs. While the association of NLR functionality and high-expression occurs broadly across plant species, this may be non-uniform across different classes of NLRs and dependent on additional mechanisms. Several of the new *Pgt* resistant NLRs fall into phylogenetic clades that also include known rust resistance genes, including *Sr35*, *Sr33*, and *Sr50*. Five of the new resistant NLRs were found in the C7 clade. Members of the C7 clade are known to function as pairs, often with members of the MIC1 (C16) clade (Bailey *et al*., 2018; Cesari, Kanzaki, *et al*., 2014; Wang *et al*., 2013). NLRs in pairs have distinct roles: the C16 NLRs are ‘sensors’ that recognise pathogen effectors and they require the C7 NLR ‘helpers’ to activate immune signalling following recognition (Cesari, Bernoux, *et al*., 2014). Here, the transfer of single C7 clade NLRs from divergent grass species provided resistance in wheat. These hits may represent a newly described function for ‘helper’ NLRs in direct pathogen recognition, or alternatively, they may be functioning with a novel partner already present in the Fielder genetic background. If so, introducing new ‘sensor’ NLRs or components of NLR pairs would provide a means to expand or revive pathogen recognition with endogenous genes (Contreras *et al*., 2022). NLR networks have previously only been described in dicot asterid plants such as *Solanaceae* species (Adachi & Kamoun, 2022; Wu *et al*., 2017), but the potential for cross-species pairings of NLRs broadens the capacity for functional resistance.

Despite their useful function in crop protection, the concept that highly expressed NLRs may be detrimental remains in the literature following early observations of deleterious phenotypes from investigations of resistance genes (Balint-Kurti, 2019; Bergelson & Purrington, 1996; Brown, 2002; Brown & Rant, 2013; Howles *et al*., 2005; Karasov *et al*., 2017; Lai & Eulgem, 2018; Richard *et al*., 2018; Stokes *et al*., 2002; Tan *et al*., 2007). The induction of NLR expression following pathogen infection similarly supported the hypothesis that all NLRs are expressed at low levels in unchallenged plants (Halterman *et al*., 2003; Tan *et al*., 2007). Few studies have directly linked NLR expression to fitness effects (Deng *et al*., 2017; Li *et al*., 2001; Oldroyd & Staskawicz, 1998; Tian *et al*., 2003; Yang *et al*., 2013; Yi & Richards, 2007; Yi & Richards, 2008). The clearest example is *A. thaliana* NLR *RPM1*, which was found to contribute to a decrease in seed production during field experiments (Tian *et al*., 2003). However, we observed the requirement of high expression for NLR function through the investigation of natural *Mla* alleles present as multiple copies in the genome, where multiple transgene inserts and high NLR expression is required for full complementation of resistance. As other *Mla* alleles directly recognise effectors from *Bh* (Saur *et al*., 2019), increased gene expression is potentially an evolutionary mechanism to compensate for any reduction in binding affinity towards their recognised effector. This would ensure threshold requirements for disease resistance under a positive dosage model (Innan & Kondrashov, 2010). In this study, no obvious macroscopic growth detriments were observed across transgenic wheat lines carrying highly expressed NLRs in the greenhouse or field experiments. Moreover, transgenic lines were obtained for 99.6% of all transformed NLRs. Further multi-year and environment testing in the absence of disease is required to detect small differences in agro-morphological traits, but the absence of pleiotropic detrimental phenotypes is consistent with the cross-species transfer of NLRs into barley (Hatta *et al*., 2021) and a multi-transgene-cassette transformed into wheat (Luo *et al*., 2021).

Given the observed trend of high expression and absence of significant pleiotropic developmental effects in recent examples, why have some NLRs caused deleterious phenotypes? New understanding of NLR function can alleviate some of these detrimental effects. For example: cell death or yield penalties caused by some NLRs can be suppressed via co-expression with their required partner or suppressor, as for *RGA4*/*RGA5* (Cesari, Bernoux, *et al*., 2014; Cesari, Kanzaki, *et al*., 2014)*, PigmR/PigmS* (Deng *et al*., 2017), and *RPS4/RRS1-R* (Guo *et al*., 2021; Zhang *et al*., 2004). Other deleterious effects may be caused by mutations in the sequence or regulatory requirements of NLRs rather than increased expression levels. In *A. thaliana*, overexpression of the wild-type *SSI4* TIR-NLR sequence did not cause the stunting and cell death observed in the mutant variant, indicating the phenotypes were caused by sequence mutation inducing auto-activity and not expression level (Shirano *et al*., 2002). Similarly, overexpression of *RPW8* alleles under the 35S promoter showed no spontaneous cell death, whereas cell death was correlated with increased transgene copies of the genomic fragment carrying *RPW8* under the native promoter (Xiao *et al*., 2003). This suggests the disruption of gene regulation as causal, such as by WRKY transcription factors (X.-M. Yang *et al*., 2024). The reduced height observed from induced mutations in *Rht13b* in wheat were also independent of transgene copy number or gene expression (Borrill *et al*., 2022). Described examples of deleterious NLRs also predominantly involve those that monitor the molecular status of host proteins or contain additional integrated host protein domains (Tian *et al*., 2003; Tong *et al*., 2017). *RPM1* is lowly expressed in transcripts in *A. thaliana* Col-0, as are other known NLRs that guard host proteins such as *LOV1* (Lorang *et al*., 2012; Tian *et al*., 2003). Different expression levels are observed across ecotypes for alleles, so effects may also be genotype dependent. In comparison, the expression of NLRs that function as singletons and those that are hypothesised to directly recognise pathogen products may not perturb host processes to the same degree. Certainly, a high expression threshold of *Mla7* is required for sufficient resistance, and several *Mla* alleles directly recognise *Bh* AVR_a_ effectors (Saur *et al*., 2019). A sufficient abundance of protein may also be required to form NLR resistosome structures (Wang, Hu, *et al*., 2019), especially for ‘helper’ NLRs (Ahn *et al*., 2023; Contreras *et al*., 2023; Ma *et al*., 2024). These results, alongside further advances in our understanding of NLR function, may explain the observation of deleterious phenotypes via mechanisms that are independent of NLR expression.

The utility of our pipeline is as an enrichment tool, as further studies will be needed to fully elucidate the biological mechanisms and function of NLR expression. Our use of a constitutive promoter for NLR screening has enabled the identification of new functional resistance genes. Subsequent work is required for optimisation; for example, NLRs in the array not conferring resistance may need additional regulatory or promoter elements for function so it is not possible to estimate the false negative rate. Alternatively, these NLRs may provide resistance to other pathogens. Testing increased expression of lowly expressed NLRs would distinguish between the effects of expression level and NLR sequence for function, and further work should consider the disruption of endogenous gene regulation. When expressed under a constitutive promoter in transient assays, lowly expressed *Rpi-amr1* transcripts were non-functional for resistance against *Phytophthora*, their presence in stable transgenic lines may cause additional effects. Are these susceptible transcripts ineffective, are they required for gene regulation, or do they confer novel recognition? How do new NLRs avoid autoactivation and unintended effects? NLRs are under strong evolutionary pressure that often leads to sequence diversification and expansion of pathogen recognition specificities (Meyers *et al*., 2005; Michelmore & Meyers, 1998; Tamborski & Krasileva, 2020; Van de Weyer *et al*., 2019). Mutation and genetic recombination of NLRs can result in the mis-regulation of immune responses, developmental effects, hybrid necrosis, and complete plant death (Ariga *et al*., 2017; Borrill *et al*., 2022; Chae *et al*., 2014; Shirano *et al*., 2002; Tran *et al*., 2017; van Wersch *et al*., 2016; Wang & Balint-Kurti, 2015). To prevent this, NLRs may initially be repressed, and only de-repressed upon providing effective resistance against pathogens without deleterious effects (Shivaprasad *et al*., 2012; Zhang *et al*., 2016). This de-repression model would predict that expressed functional NLRs will be present in genomic regions with reduced methylation and open chromatin structure. Furthermore, in this model NLRs that are highly expressed have a higher likelihood of being functional without deleterious effects.

By exploiting a simple observation of the association between functionality and high expression, we have produced a high-throughput pipeline, designated NLRseek™, which facilitates large-scale mining of resistance genes from a wide and diverse gene pool. This approach provides a path to access genetic resistance in diverse plant species previously inaccessible to traditional methods. High-throughput phenotypic screening of plant transgenic arrays has been used successfully for identifying drought tolerance in rice (Komori *et al*., 2020) and is a powerful tool for trait discovery in crops. The use of transgenic arrays enables the study of pathogens less amenable to *in vitro* culture and molecular investigation, such as shown here for *Pgt*. In addition, NLR arrays can also be validated through transient transformation, protoplast-based assays, and other methods of high-throughput screening. The many diverse NLRs found using this approach support a breadth of variation to deploy tailored gene stacks against major pathogens (Hafeez *et al*., 2021; Luo *et al*., 2021). Moreover, AI tools can enable greater insights into structure-function associations, paving the way for engineering to produce new resistance genes for emerging disease threats. The signature of high expression, combined with advances in gene cloning and genotype-independent transformation technology, ignites an exciting potential for the future of biotechnology to protect plant health and improve food security.

## Methods

### Native copy number variation of *Mla7*

Genomic DNA was extracted using a CTAB extraction approach from the Manchuria near-isogenic lines CI 16147 and CI 16153, which carry *Mla7* from two different donors (Multan and Long Glumes) (Moseman, 1972). Illumina sequencing was performed at Novogene (Cambridge, UK) using 150 bp paired end reads. Reads were trimmed using Trimmomatic (v0.39) with adapter clipping using TruSeq3-PE adapters and parameters 2:30:10, trimming of low quality leading and trailing sequence with parameter 5, sliding window trimming with parameters 4:10, and a final minimum length of 36 bp. Copy number of *Mla7* and the control gene *Bpm* was performed using the k-mer analysis toolkit (KAT). The module sect was used with default parameters to determine k-mer coverage over the gene sequences of *Mla7* and *Bpm*. R (v4.1.2) and ggplot2 (v3.3.6) were used to estimate copy number variation of *Mla7*.

### Transgenic complementation of *Mla7*

For assembly of the *Mla7* construct, the native genomic fragment of *Mla7* and promoter, UTRs, and terminator regions of *Mla6* were amplified from barley accessions CI 16153 (*Mla7*) and CI 16151 (*Mla6*) (Halterman *et al*., 2001), using GoTaq Long PCR Master Mix (Promega). Construct development was performed as described in Bettgenhaeuser *et al.,* (2021). Briefly, PCR fragments were assembled into pBract202 binary vector (BRACT) via Gibson reaction (Gibson, 2014). Primers for PCR fragments bearing 20 bp overlapping to the flanking region were produced with Phusion High-Fidelity DNA Polymerase (NEB) and 40-mer primers. Barley transformation was performed using the technique described by Hensel *et al*. (2009) using the hygromycin resistance gene (*hyg*) as a selectable marker. The barley powdery mildew (*Bh*) and wheat stripe rust (*Pst*) susceptible SusPtrit x Golden Promise DH-47 from the SxGP doubled haploid population (Yeo *et al*.) was used as the transformable background. Copy number variation in transgenic plants was determined by quantitative real time PCR using the selectable marker gene *hyg* similar to the approach by Bartlett *et al*. (2008) and was performed by iDna Genetics (Norwich, UK). A population segregating for two T-DNA inserts was generating by crossing transgenic lines T1-117 and T1-121 (SxGP DH-47 transformed with *pMla7*::*Mla7::tMla7*) with the *Bh* and *Pst* susceptible accession Manchuria. T-DNA were mapped in segregating F_2_ populations to chromosomes 3H and 5H, respectively, using SNP markers derived from the OPA markers 1_0702 and 2_1012 (Close *et al*., 2009). F_2_ lines heterozygous for the T-DNA and homozygous at the *Mla* locus for the Manchuria allele (non-functional *mla* allele) were crossed and validated in the resulting progeny for its allelic state at both T-DNA and *Mla* loci. Selfed seed of this accession was used for pathogen assays with *Bh* and *Pst*.

### Pathogen assays for *Mla7*

Pathogen assays with *Bh* and *Pst* were performed as described in Bettgenhaeuser *et al.,* (2021). A collection of *Bh* isolates (N=13) were selected from a gene bank of the pathogen containing 59 reference isolates collected in 12 countries in all nonpolar continents over a period of 66 years (1953 – 2019) and kept at the Agricultural Research Institute Kroměříž Ltd. Virulence patterns to 35 standard barley genotypes are shown in Dreiseitl (2019). Prior to inoculation, isolates were checked for their purity and their correct virulence phenotypes were verified on standard barley lines (Kølster *et al*., 1986). *Bh* isolates were multiplied on leaf segments of a susceptible cultivar Bowman. *Bh* isolate CC148 was propagated on barley cv. Manchuria (CI 2330) prior to inoculation. Seedlings were placed horizontally, inoculated, rotated 180°C after a resting period of 2 min for conidia to settle, and inoculated again. Susceptible controls in every experiment included Manchuria (CIho 2330), Pallas (CIho 11313), Siri (CIho 14846), and the barley cv. Siri-derived set of near-isogenic lines, each carrying a single mildew resistance gene (Kølster & Stølen, 1987). *Bh* isolate CC148 assays were carried out in a negative pressure containment greenhouse with supplemental lighting and temperature set at 18°C day and 12°C night. For pathogen assays using the diverse collection of *Bh* isolates, inoculation and evaluation protocols were recently described in detail (Dreiseitl, 2022). Briefly, seed was grown in a mildew-proof greenhouse under natural daylight. Central leaf segments of 15mm were cut from fully expanded primary leaves after 14 days for each transgenic family and controls. Leaves were placed on water agar (0.8%) containing benzimidazole (40 mg/L) in Petri dishes with adaxial surfaces facing upwards. A settling tower was used for inoculation: leaf segments were placed at the bottom of the tower and conidia from a fresh leaf segment of the susceptible cultivar were blown in the settling tower at a concentration of approximately 8 conidia/mm^2^. The Petri dishes of leaf segments were incubated at 20 ± 2°C under artificial light (cool-white fluorescent lamps providing 12 h light at 30 ± 5 μmol m^-2^ s^-1^). Infection responses were scored seven days after inoculation on a 0-4 scale where: 0 = no visible mycelium or sporulation, and 4 = strong mycelial growth and sporulation (Torp *et al*., 1978). Scoring was repeated a day later and two replications were performed.

Briefly, *Pst* inoculations were performed using a suspension of urediniospores in talcum powder, at a weighted ratio of spores:talcum powder of 1:16 and applied to leaves using a spinning table. Plants were sealed and stored at 8 °C for 48 h immediately after inoculation. Plants were grown prior to and after inoculation in a controlled environment room under 16h light/8 hr dark. Phenotypes of the first leaves were scored 14 days post-inoculation using on a 0.5 incremental scale of 0 to 4 representing the surface area displaying an infection phenotype where: 0 represented no chlorosis or no pustules of *Pst*, and a score of 4 indicated infection across 100% of the surface area.

### Plant materials and growth conditions for RNAseq analysis

Seeds of the following grass species were used for the preparation of libraries of candidate NLR genes: *Achnatherum hymenoides*, *Aegilops bicornis*, *Aegilops longissima*, *Aegilops searsii*, *Aegilops sharonensis*, *Agropyron cristatum*, *Avena abyssinica*, *Brachypodium distachyon*, *Briza media*, *Cynosurus cristatus*, *Echinaria capitata*, *Holcus lanatus*, *Hordeum vulgare*, *Koeleria macrantha*, *Lolium perenne*, *Melica ciliata*, *Phalaris coerulescens*, *and Poa trivialis*.

Seeds were germinated on damp filter paper on petri dishes and placed at 4°C for 6-7 days to break seed dormancy. Germinated seeds were transferred into an in-house custom soil mix prepared by the horticultural services department at The John Innes Centre (JIC cereal mix: 65% peat, 25% loam, 10% grit, 3 kg/m^3^ dolomitic limestone, 1.3 kg/m^3^ PG mix, and 3 kg/m^3^ Osmocote Exact). Seedlings were grown in a clean controlled environment chamber under 16 hrs light at 20°C/8 hrs dark at 16°C. The controlled environment chamber used is a clean germination room free of pests and disease. For RNA isolation, leaves of plants 12 to 35 days post germination were used depending on the species. The first and second leaves of plants were harvested for most plant species; however, the first to sixth leaves were used from species with smaller leaf sizes.

Seeds of *Solanum lycopersicum* cultivars VFNT Cherry (LA1221) and Motelle (LA2823) were obtained from The C.M. Rick Tomato Genetics Resource Center (TGRC; https://tgrc.ucdavis.edu/). Seeds were germinated in an in-house custom soil mix prepared by the horticultural services department at The John Innes Centre (JIC multipurpose + grit: 90% peat, 10% grit, 4kg/m^3^ dolomitic limestone, 0.75 kg/m^3^ PG mix, and 1.5 kg/m^3^ Osmocote Bloom). Seedlings grown in a clean controlled environmental chamber free from pests and disease under 16 hrs light/8 hrs dark at 18°C. Tissue was sampled for the RNA isolation after one month. Fully expanded leaves were used for leaf tissue and the entire root system was used after washing in distilled water. For each tissue type, samples were pooled from three seedlings per cultivar.

For *Arabidopsis thaliana*, seeds of the lines Col-0, Ler-0, Mt-0, and Ws-0 were obtained from the Nottingham Arabidopsis Stock Centre (https://arabidopsis.info/). Seeds were surface sterilised in a sterilisation chamber using chlorine gas for 5 hours and sown on Murashige and Skoog (MS) media with 1% sucrose supplemented with 0.8% agar. Seeds were kept at 4°C for 2-3 days and then grown at 22°C under 16 hr light. Seedlings were transferred to liquid MS media with 1% sucrose in a 24 well tissue culture plate with seedlings in each well. The controlled environment chamber used is a clean germination room free of pests and disease. Seedlings were sampled for RNA isolation after 9-10 days post-germination.

### RNA extraction and sequencing

Total RNA was extracted from leaves of grass species, seedlings of *A. thaliana*, and leaf and root tissue of *S. lycopersicum* using a Trizol-phenol based protocol according to manufacturer’s protocol (Sigma-Aldrich; T9424). Barcoded Illumina TruSeq RNA HT libraries were constructed and pooled with four samples per lane on a single HiSeq 2500 lane run in Rapid Run mode. Sequencing was performed using 150 bp paired-end reads. Paired-end reads were assessed for quality using FastQC and trimmed before assembly using Trimmomatic (v0.36) with parameters set at ILLUMINACLIP:2:30:10, LEADING:5, TRAILING:5, SLIDINGWINDOW:4:15, and MINLEN:36. These parameters were used to remove all reads with adapter sequence, ambiguous bases, or a substantial reduction in read quality. De novo transcriptome assemblies were generated using Trinity with default parameters (version 2013-11-10). kallisto (v0.43.1) was used to estimate expression levels for all transcripts using default parameters and 100 bootstraps.

### *Rpi-amr1* isoform characterisation and *Phytophthora infestans* assays

Isoforms were identified from the transcriptome of *Solanum americanum* accession SP2273 using BLAST (v2.2.31) of *Rpi-amr1* (GenBank: MW348763, NCBI). Sequence analysis performed using Geneious Prime (v2024.0.3). Gene isoforms were synthesised via Gene Universal due to inability to amplify lowly expressed isoforms from cDNA. CDS sequences were expressed under the NRC4 promoter and terminator and constructs were assembled using Golden Gate into the Level 1 acceptor pICH47732. Constructs of *Rpi-amr1* from resistant accession SP2273, *Rpi-amr1* from susceptible accession SP2271, and *Rpi-amr3* from SP1102 published in Witek *et al*. (2021) used as controls. Transient complementation assays and *P. infestans* inoculation were performed as described previously (Foster *et al*., 2009; Witek *et al*., 2016). Briefly, *Agrobacterium* liquid cultures were re-suspended in MES buffer (10 mM MES, 10 mM MgCl2, 150 mM acetosyringone) and adjusted to 0.3 OD_600_. A total of 3 to 4 leaves of different *N. benthamiana* were infiltrated with each construct per replicate with a total of 3 replicates performed. *P. infestans* 88069 was grown on rye media and sporangia harvested after 10 days. Leaves were inoculated with two 10-μl droplets of a zoospore suspension (50,000 zoospores ml^-1^). Inoculated leaves were incubated for 7 to 12 days on damp paper towels in a Sanyo cabinet at 16°C under 16hr light and 8 hr darkness before phenotypes scored. The lesion area was measured from images using Fiji (ImageJ2, v2.14.0/1.5f) and analysed using RStudio (2023.12.1+402).

### Identification of highly expressed NLRs

TransDecoder (v4.1.0) LongOrfs was used to predict all open reading frames in *de novo* assembled transcriptomes. Transcript abundance was quantified using kallisto (Bray *et al*., 2016). InterProScan (v5.27-66.0) was used to annotate domains using Coils and the Pfam, Superfamily, and ProSite databases. Any protein that contained both a nucleotide-binding domain and a leucine-rich repeat domain was advanced in the analysis. Histograms were generated using RStudio (v2023.12.1+402). The transcripts of known and characterised NLRs were identified from the transcriptome using a BLAST+ (v2.2.31) search using the publicly available nucleotide sequence. Sequence similarity of the CDS was assessed using MUSCLE (v5.1) using default parameters.

### Building the NLR array and molecular cloning

Sequencing, *de novo* RNAseq assembly, NLR identification, and PCR primer development were completed for 81 plant accessions from 18 grass species. Of the 81 accessions sequenced, 69 resistant accessions were progressed to molecular cloning including species in the genera *Achnatherum, Aegilops, Agropyron*, *Avena, Brachypodium, Briza, Cynosurus, Echinaria, Holcus, Hordeum*, *Koeleria*, *Lolium, Melica, Phalaris,* and *Poa*.

Highly expressed NLRs were identified according to pipeline described above, with the addition of Fast Approximate Tree Classification (FAT-CAT; Afrasiabi *et al*., 2013) which was used to classify nucleotide-binding domains based on a phylogenetic tree developed from rice, *Brachypodium distachyon*, and barley nucleotide-binding domains derived from NLRs (Bailey *et al*., 2018). NLR encoding genes were advanced based on the following requirements: the transcript must contain either a complete or 5’ partial open reading frame; the gene must be among the top 25% expressed NLRs; and the gene does not belong to NLR families known to require an additional NLR (Bailey *et al*., 2018). Among the candidate NLRs, redundancy was removed using CD-HIT (v4.7) requiring 100% identity (-c 1.0).

For molecular cloning, PCR primers were developed using Gateway adapters attB1 and attB2 fused to first 20 nucleotides of the start or end of the coding sequence, respectively. The proportion of cloned NLRs is variable according to species, guided by the available diversity of accessions in each species and the prevalence of resistance to target pathogens. PCR primers were developed for a total of 1,909 NLRs. In total, 1,019 NLRs were cloned into Gateway pDONR entry vector. This set includes known resistance genes: *Sr33* (wheat stem rust; (Periyannan *et al*., 2013), *Sr35* (wheat stem rust; (Saintenac *et al*., 2013), and *Sr50* (wheat stem rust; (Mago *et al*., 2015), *Lr21* (wheat leaf rust; Huang *et al*., 2003), *Yr10* (wheat stripe rust; (Liu *et al*., 2014)), *Pm3b* (wheat powdery mildew; (Srichumpa *et al*., 2005; Yahiaoui *et al*., 2004), *Mla3* (barley powdery mildew and rice blast (Brabham *et al*., 2024), *Mla7* (barley powdery mildew) and *Mla8/Rps7* (barley powdery mildew and wheat stripe rust; (Bettgenhaeuser *et al*., 2021; Halterman & Wise, 2004).

### Plant transformation

The NLRs in the entry clones were transferred to a destination vector pDEST2BL, which is a binary vector, by LR reaction of the Gateway® system. NLRs were expressed under the maize ubiquitin promoter. The destination vector pDEST2BL includes the DsRed2 fluorescent protein used as a visual selectable marker in the seed (Yarbrough *et al*., 2001). The resultant transformation vectors were introduced into *Agrobacterium tumefaciens* strain EHA105 by electroporation. The *Agrobacterium* strains carrying the transformation vectors were used to transform wheat cv. Fielder according to the published method (Ishida *et al*., 2015) with a modification that an immature embryo was cut into three pieces when transferred to the second selection medium. Fielder seeds were obtained from Kihara Institute for Biological Research, Yokohama City University. Briefly, 15 immature embryos were infected with each of the *Agrobacterium* strains and up to seven independent events per NLR were grown to maturity. A total of 6,260 transformation events were achieved for 999 NLRs and seed was obtained from T_1_ plants from 995 NLRs. Fifty or more seeds were obtained from 96.4% of the harvested events. A total of 5,177 T_1_ families were generated, families were further subdivided based on the fluorescence of the seed to a total of 10,646 DsRed2 groups.

### Inoculation and phenotyping with the wheat stem rust pathogen (*Puccinia graminis* f. sp. *tritici*) and the wheat leaf rust pathogen (*Puccinia triticina*) at the seedling stage in the greenhouse

The independent T_1_ families within an NLR construct were subsampled and three seedlings from each T_1_ family were used for resistance phenotyping. Rust inoculations were made according to the standard protocols used at the USDA-ARS Cereal Disease Laboratory and the University of Minnesota (Huang *et al*., 2018). Briefly, on the day before inoculation, urediniospores of the rust pathogens were removed from a –80°C freezer, heat-shocked in a 45°C water bath for 15 min, and then rehydrated in an 80% relative humidity chamber overnight. After assessing the germination rate (Scott *et al*., 2014), 10 mg of urediniospores were placed into individual gelatin capsules (size 00) to which 700 ml of the light mineral oil (Soltrol 170, Chevron Phillips Chemical Company) carrier was added. The inoculum suspension was applied to 12-day-old plants (second leaf fully expanded) using custom atomizers (Tallgrass Solutions, Inc., Manhattan, KS) pressured by a pump set at 25 to 30 kPa. Approximately 0.15 mg of urediniospores were applied per plant. Immediately after inoculation, the plants were placed in front of a small electric fan for 3 to 5 min to hasten evaporation of the oil carrier from leaf surfaces. Plants were allowed to off-gas for an additional 90 min before placing them inside mist chambers. Inside the mist chambers, ultrasonic humidifiers (Vick’s model V5100NSJUV; Proctor & Gamble Co., Cincinnati, OH) were run continuously for 30 min to provide sufficient initial moisture on the plants for the germination of urediniospores. For the next 16 to 20 h, plants were kept in the dark and the humidifiers run for 2 min every 15 min to maintain moisture on the plants. Light (400W high pressure sodium lamps emitting 300mmol photon s−1m−2) was provided for 2 to 4 h after the dark period. Then, the chamber doors were opened halfway to allow the leaf surfaces to dry completely before returning the plants to the greenhouse under the same conditions described above (Huang *et al*., 2018).

All rust phenotyping experiments were conducted in a completely randomized design. Accessions exhibiting variable reactions across experiments were repeated in an additional experiment if sufficient seeds were available. Stem rust infection types (ITs) on the accessions were scored 12 days after inoculation using a 0 to 4 scale (Roelfs & Martens, 1988; Stakman *et al*., 1962). Original seedling IT data were converted to a numerical 0-9 linear scale as detailed in Zhang *et al*., 2014. The initial phenotyping of transgenic lines was performed with the Minnesota, USA *Pgt* race QTHJC (isolate 69MN399 provided courtesy of Yue Jin, USDA-ARS Cereal Disease Laboratory, St. Paul, MN USA). Lines found resistant to *Pgt* race QTHJC were further phenotyped against the more broadly virulent *Pgt* race TTKSK (isolate 04KEN156/04) from Kenya. Both races are virulent on the wheat cultivar Fielder. Seed of T_2_ individuals was used for photographs of QTHJC infection and for NLRs NLR_05_75, NLR_05_92, NLR_08_16, NLR_08_79, and NLR_09_55 for TTKSK screening due to seed availability.

Wheat leaf rust screening was carried out using a protocol similar to that used for wheat stem rust screening but without the light period being provided at the end of the infection period (Huang *et al*., 2018). Race THBJ (isolate 99 ND 588 provided by James Kolmer, USDA-ARS Cereal Disease Laboratory, St. Paul, MN USA) was used for the assays. Leaf rust ITs were scored after inoculation using a 0 to 4 scale (Kolmer, 2013) and data were converted to a numerical 0-9 linear scale as described above.

### Inoculation and phenotyping with the wheat stem rust pathogen (*Puccinia graminis* f. sp. *tritici*) at the adult stage in the field

Individual lines found resistant in the seedling assays with *Pgt* race QTHJC were bulked and T_2_ families from multi-transgene lines and control lines were planted at the University of Minnesota Rosemount Research and Outreach Center in Rosemount, MN in 2021 and 2022. The wheat cultivars Fielder, Morocco, and LMPG-6 were used as susceptible controls. The same isolate of *Pgt* race QTHJC used in the seedling assays was also used for the adult plant assays in the field. Field inoculations were performed as previously described in Luo *et al*., 2021. Several of the lines found resistant in the seedling stage were not available for testing in the field due to low seed production and therefore partially overlapping field trials were used.

Approximately 25 seeds per line were planted in each plot. When the first nodes of plants were detectable (31 Zadoks scale), they were inoculated with a suspension of *Pgt* urediniospores (1 g urediniospores per 1 L Soltrol 170 mineral oil; Phillips Petroleum) using an ultra-low volume sprayer (Mini-ULVA, Micron Group). Three additional inoculations were made in successive weeks to ensure high infection levels during the later stages of crop development. Severity was recorded as the visual percentage (0-100%) of stem and leaf sheath tissue covered by uredinia. Ratings were assessed using the modified Cobb scale (Peterson *et al*., 1948). The infection responses (IRs) were recorded as highly resistant (HR; clear hypersensitive infection sites but with no pathogen sporulation), resistant (R; minute-to-small uredinia surrounded by chlorosis or necrosis), moderately resistant (MR; medium-sized uredinia often surrounded by chlorosis), moderately susceptible (MS; medium-to-large erumpent uredinia with little or no chlorosis), or susceptible (S; very large erumpent uredinia with little or no chlorosis).

### Phylogenetic analyses

Phylogenetic analysis of the NB domains of NLRs was carried out as described by Bailey *et al*., (2018) using updated NLR gene annotations for barley (Mascher *et al*., 2021), *B. distachyon* (v3.1; NCBI PRJNA32607 and PRJNA74771), and wheat (Zhu *et al*., 2021). NB domains from NLRs were identified using HMMer (v3.3.2) hmmalign with the hidden Markov model encompassing the NB-ARC1-ARC2 domains (Bailey *et al*. 2018). The alignment was converted to an aligned FASTA file using esl-reformat and processed using the QKphylogeny script QKphylogeny_alignment_analysis.py with parameters ‘-d 0.3’ (non-redundant), ‘-b 0.5’ (breadth coverage of alignment greater than or equal to 50%), and ‘-d 0.3’ (depth of coverage at each residue of greater than or equal to 30%). The phylogenetic tree was constructed using RAxML (v8.2.12) using the PROTGAMMAJTT model and 1,000 bootstraps and visualized using iTOL (https://itol.embl.de/).

The Pooideae species phylogenetic tree was generated using the QKbusco pipeline (https://github.com/matthewmoscou/QKbusco). BUSCO (v3.0.2) with default parameters and embryophyte_odb9 library was used to identify genes using annotated coding sequences (sequenced genomes) or open reading frames predicted using TranDecoder (v4.1.0) from *de novo* assemblies (transcriptomes). QKbusco_merge.py was used to parse BUSCO output and prepare FASTA files for multiple sequence alignment. The parameter status was set to fragmented to allow fragmented coding sequence to be included in the analysis. Codon-based multiple sequence alignment of individual genes was performed using PRANK (v.170427). Individual gene multiple sequence alignments were merged using QKbusco_phylogeny.py using a coverage depth of 40% at individual sites for inclusion in the alignment. The maximum likelihood phylogenetic tree was generated using RAxML (v8.2.12) with the GTRGAMMA model.

## Author contributions

Conceptualization: MJM, HPvE, NT, RF

Data curation: MJM, HJB, BS, OM, IHP, TKomo

Formal Analysis: HJB, MJM, OM, BS

Funding acquisition: HPvE, NT, RF

Investigation: HJB, MJM, KW, IHP, AD

Methodology: MJM, HJB, IHP, CY, NI, NT, TKoma, TKomo, HN, PG, AD, AH, BS, OM, AF

Project administration: MJM, HJB, TKomo

Resources: MJM, AH, NT, BS

Software: MJM

Supervision: MJM, HJB, HPvE, BS, TKomo

Validation: MJM, HJB, KW

Visualization: MJM, HJB

Writing – original draft: HJB

Writing – review & editing: MJM, HJB, HPvE, KW, BS, OM, TKomo, TKoma, AD

## Data availability

Whole genome sequencing data of barley accessions CI 16147 and CI 16153 were deposited in NCBI BioProject PRJNA952654. RNAseq data for *Arabidopsis thaliana*, tomato, and diverse Poodieae species were deposited in NCBI BioProject PRJNA928100, PRJNA927036, and PRJNA913397, respectively. GenBank identifiers for transformation construct sequence for *Mla7* under *Mla6* promoter/terminator and native sequence are MZ555770 and OQ859100, respectively.

## Acknowledgements

The authors would like to thank Vera Sham, Mikhaela Neequaye, Henry Jennings, Sophie Sharpe, Inesh Amarnath, and Sara Perkins for their technical help. The authors would also like to thank Paul Nicholson, Yogesh Gupta, Jonathan Jones, Jack Rhodes, and Robert Heal for scientific discussion. In addition, the authors would also like to thank Diana Horvath for scientific discussion and editing of the manuscript.

## Supplemental

**Supplemental Figure 1.**
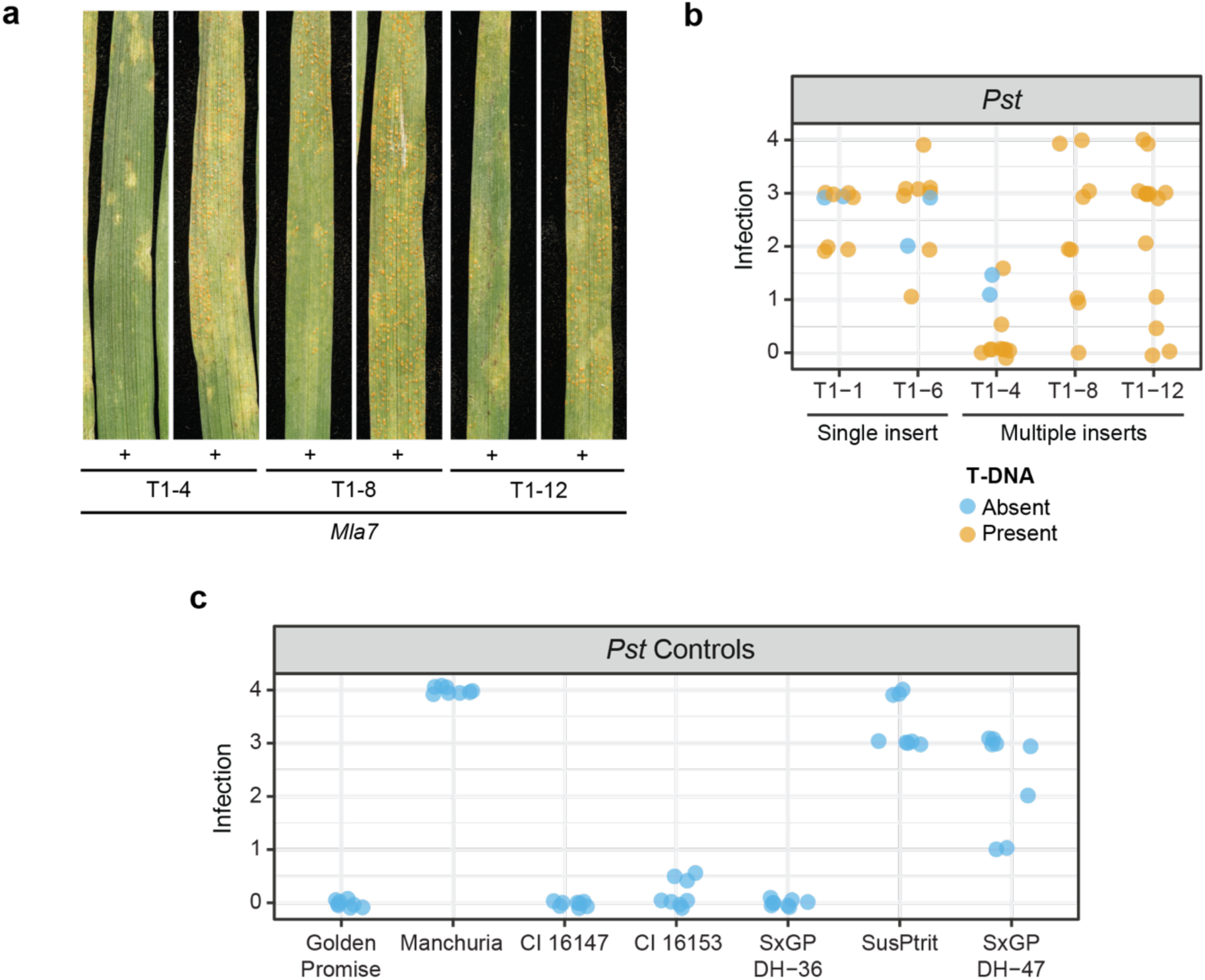
M*l*a7 confers resistance to wheat stripe rust (*P*. *striiformis* f. sp. *tritici*) isolate 16/035. Wheat stripe rust susceptible barley cv. SxGP DH-47 was transformed with *Mla7* driven by the *Mla6* promoter and *Mla6* terminator. Two single copy insert lines (T1-1 and T1-6) and three multiple copy insert lines (T1-4, T1-8, and T1-12) were identified for *Mla7*. **A)** Resistance to wheat stripe rust isolate 16/035 was observed in transgenic barley lines carrying *Mla7*. **B)** Multiple T-DNA insertions are required to recapitulate wild-type resistance to *Pst*, as only transgenic lines carrying more copies of *Mla7* displayed resistant phenotypes of 0. Infection phenotypes for individual leaves in T_1_ families. Presence or absence of T-DNA is shown in orange and blue, respectively. All phenotypes are on a scale from 0.0 to 4.0 in 0.5 increments representing the degree of leaf area covered with uredinia. Transparency and jittering were used to visualize multiple overlapping data points. **C)** Resistant control barley lines for *Pst* inoculations included Golden Promise (*Mla8*/*Rps7*), CI 16147 (*Mla7*/*Rps7*), CI 16153 (*Mla7*/*Rps7*), and SxGP DH-36 (*Mla8*/*Rps7*). Susceptible controls included Manchuria, SusPtrit, and SxGP DH-47 the genetic background used for *Mla7* transgenic lines. The experiment was carried out in parallel with results and controls of Bettgenhaeuser *et al*. (2022).

**Supplemental Figure 2.**
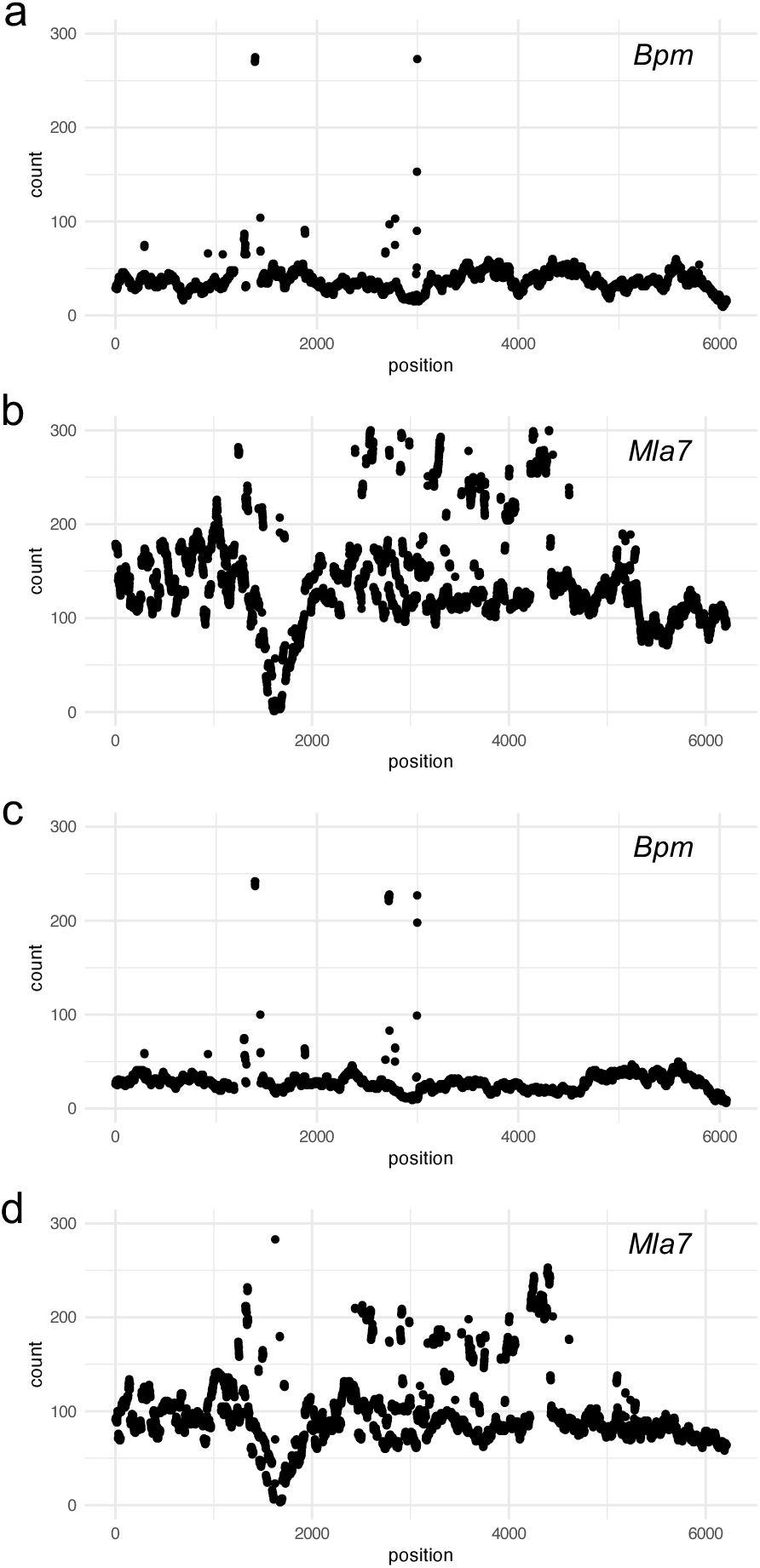
Barley accessions CI 16147 and CI 16153 carry three copies of *Mla7*. Whole genome sequencing was performed on CI 16147 and CI 16153 and k-mer analysis for the genes *Bpm* and *Mla7*. **A**) Coverage analysis of *Bpm* in CI 16147 estimates a median coverage of 36 (1 copy). **B**) Coverage analysis of *Mla7* in CI 16147 estimates a median coverage of 117 (∼3.25 copies). **C**) Coverage analysis of *Bpm* in CI 16153 estimates a median coverage of 27 (1 copy). **D**) Coverage analysis of *Mla7* in CI 16153 estimates a median coverage of 87 (∼3.22 copies).

**Supplemental Figure 3.**
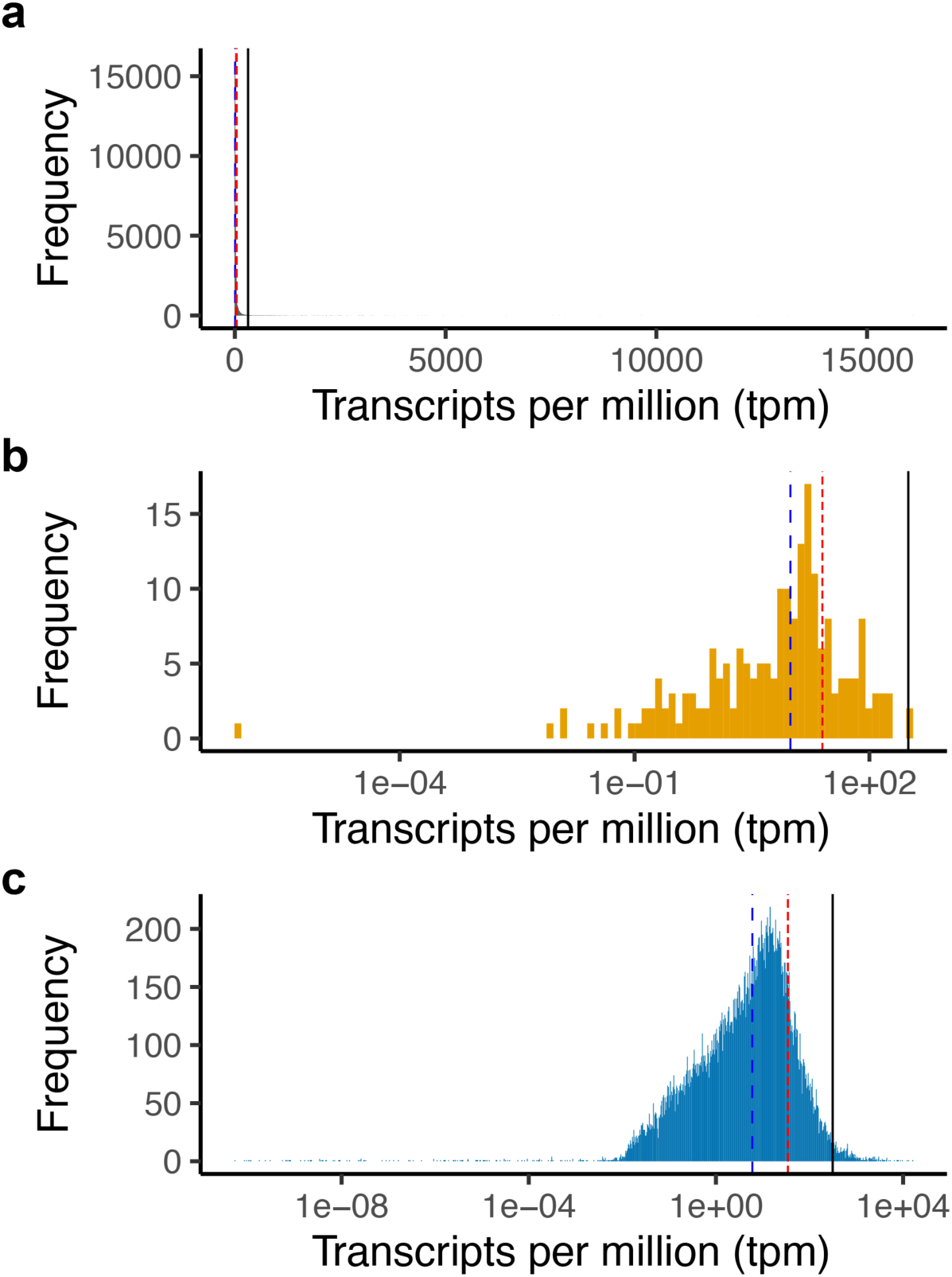
Expression level of all genes in *A. thaliana* ecotype Col-0. Median gene expression level of 5.98 tpm annotated with a blue dashed line. Mean gene expression level of 34.60 tpm annotated with a shorter dashed red line. The expression level of the highest expressed NLR at 313.68 tpm indicated with a solid black line. An average of the publicly available datasets of replicates SRR5197904, SRR5197905, and SRR5197906 used for analysis. TAIR10 gene annotations and NLR annotations from Meyers *et al*., (2003) were used. **A)** Histogram of untransformed gene expression levels. **B)** Histogram of gene expression of all NLR genes plotted on a log_10_ transformed scale of transcripts per million. **C)** Histogram of gene expression of all non-NLR genes plotted on a log_10_ transformed scale of transcripts per million.

**Supplemental Figure 4.**
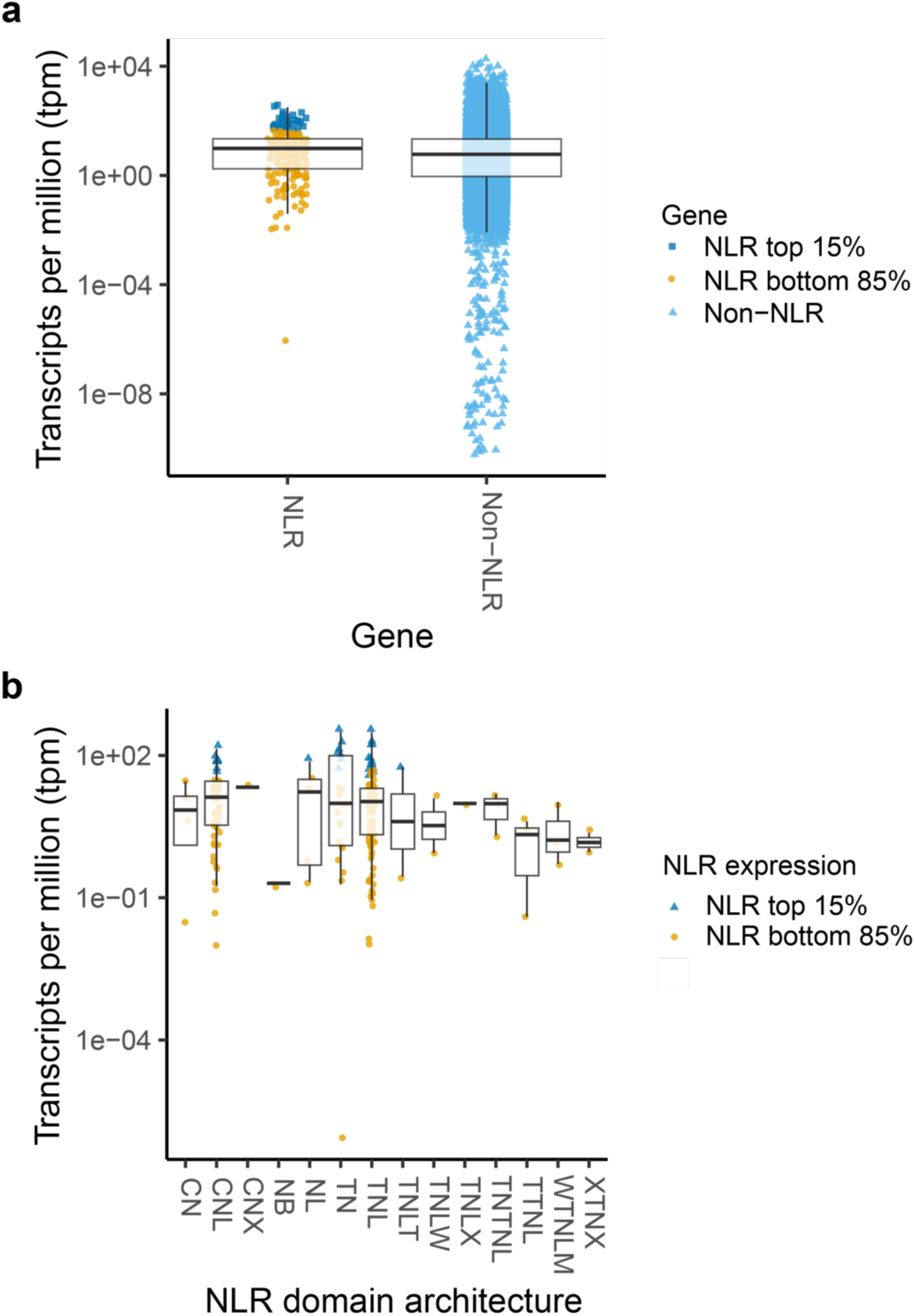
Comparative NLR expression in *A. thaliana* ecotype Col-0. Box plots of gene expression from *A. thaliana* ecotype Col-0. An average of the publicly available datasets of replicates SRR5197904, SRR5197905, and SRR5197906 used for analysis. Median gene expression indicated by the thicker horizontal line within the white box denoting the interquartile range. **A)** Expression level of NLRs compared to non-NLRs plotted on a log_10_ transformed scale of transcripts per million. The top 15% of expressed NLRs indicated with dark blue squares, the bottom 85% of expressed NLRs indicated with orange circles, and non-NLRs indicated with light blue triangles. **B)** Expression level of different NLR domain classes as classified in Meyers *et al*., (2003, 2002) plotted on a log_10_ transformed scale of transcripts per million. NLR domain architecture indicated as: CN=Coiled coil (CC)-nucleotide binding site (NBS), CNL = CC-NBS-leucine rich repeat (LRR), CNX = CC-NBS-X (where X indicates additional variable domains), NB = NBS, NL = NBS-LRR, TN = TIR-NBS, TNL = TIR-NBS-LRR, TNLT = TIR-NBS-LRR-TIR, TNLW = TIR-NBS-WRKY, TNLX = TIR-NBS-LRR-X, TNTNL = TIR-NBS-TIR-NBS-LRR, TTNL = TIR-TIR-NBS-LRR, WTNLM = WRKY-TIR-NBS-LRR-Kinase like domain, XTNX = X-TIR-NBS-X. The top 25% of expressed NLRs indicated with blue triangles and the bottom 75% of expressed NLRs indicated with orange circles.

**Supplemental Figure 5.**
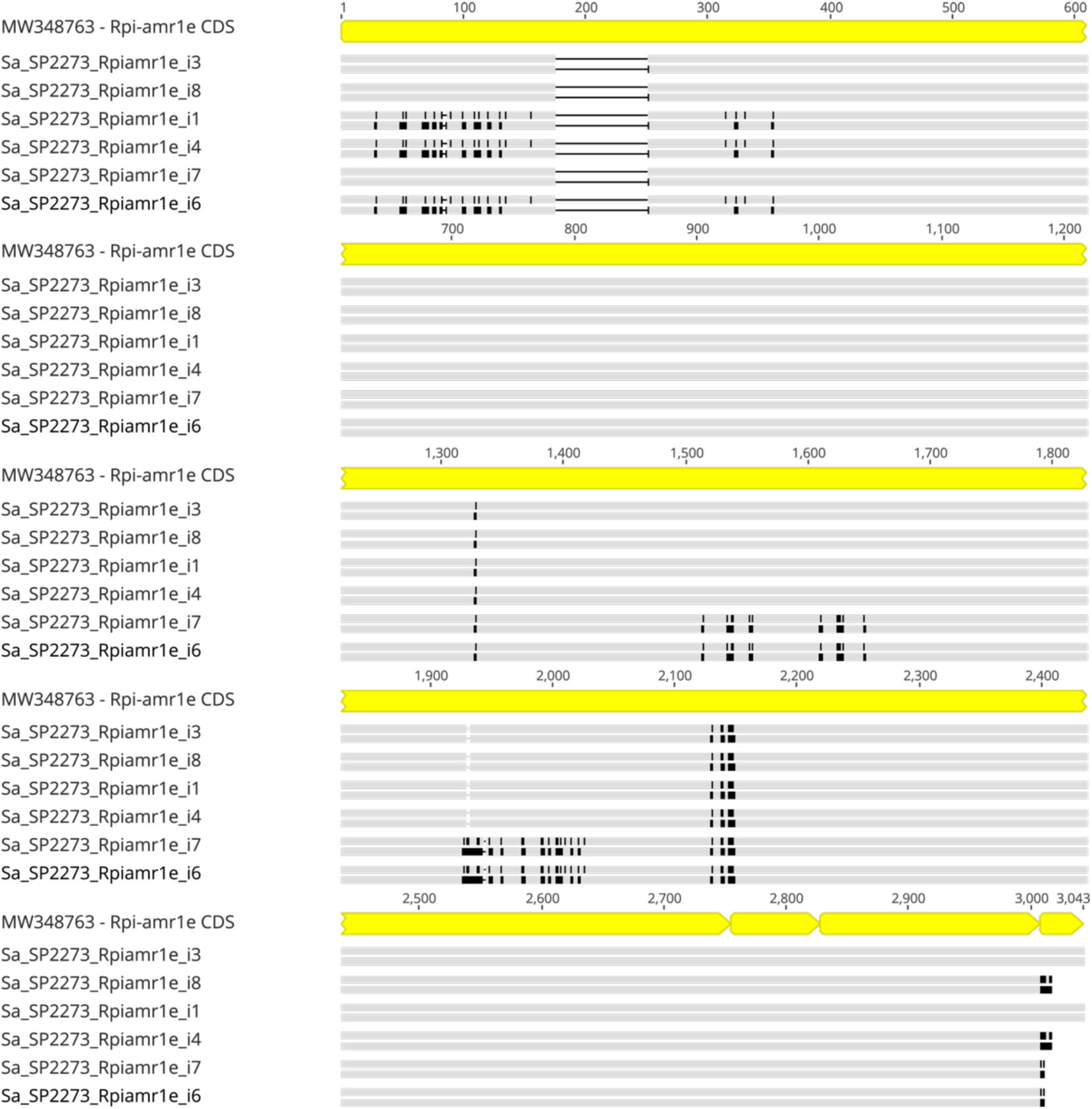
Sequence alignment of *Rpi-amr1* isoforms. Alignment of isoforms of *Rpi-amr1* identified from the *de novo* assembled transcriptome of *S. americanum* accession SP2273 against the *Rpi-amr1* publicly available sequence (GenBank: MW348763). The exon structure of *Rpi-amr1* (GenBank: MW348763) indicated in the yellow annotations. *Rpi-amr1* isoforms listed in descending order of expression level, with isoform *i3* being the most highly expressed. For each isoform, the top grey bar shows nucleotide sequence similarity, and the lower grey bar shows amino acid sequence similarity. Black bars indicate sequence differences and the black line with the break in the grey bar indicates presence/absence variation. Multiple sequence alignment performed using Geneious.

**Supplemental Figure 6.**
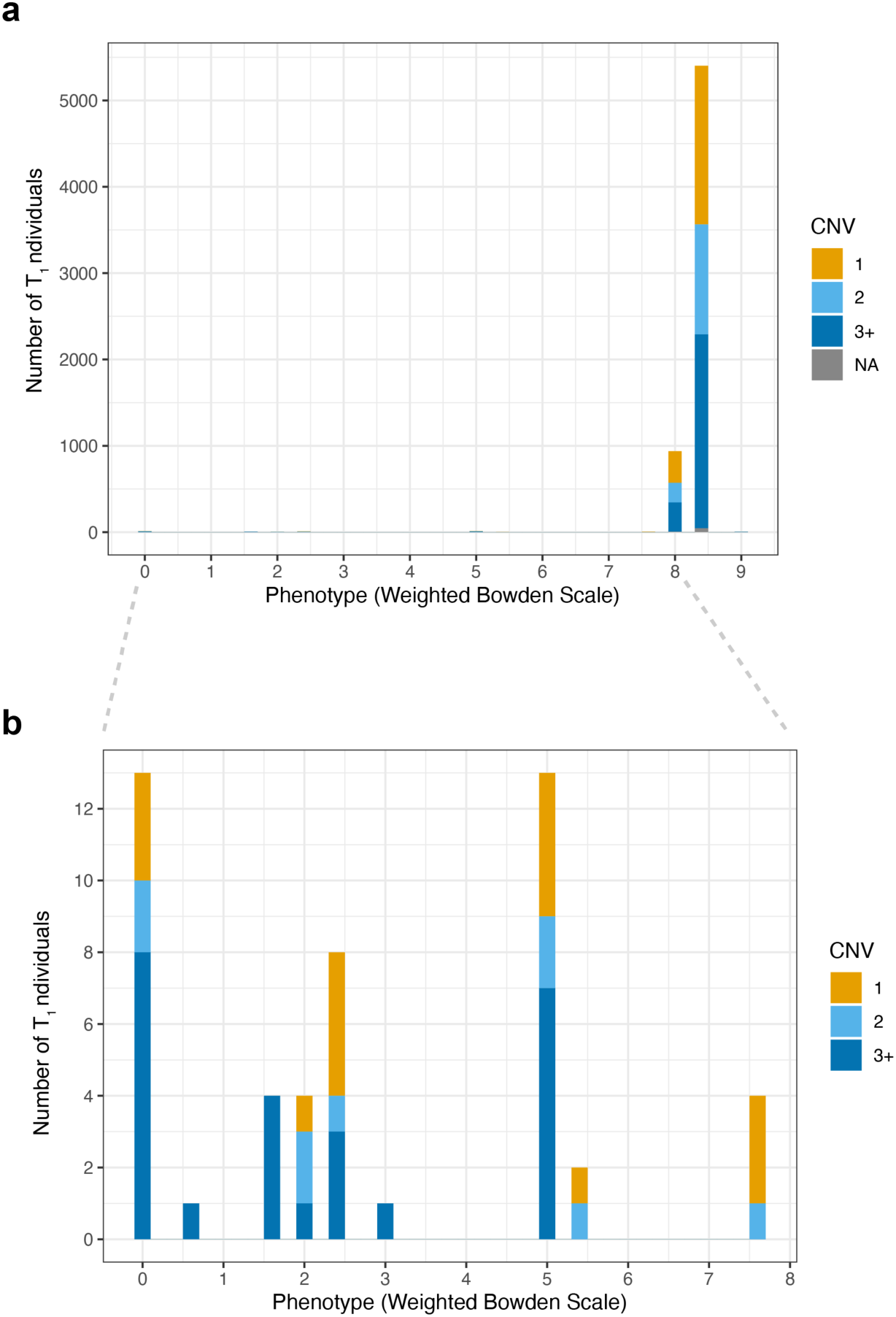
Phenotypes of individuals from the 995 NLR array following inoculation with *Puccinia graminis* f. sp. *tritici* race QTHJC. Individual phenotypic scores for each T_1_ transgenic line shown (**Supplemental Data 4**). CNV = Copy number variation. Copy number of individual T_1_ lines indicated as 1 (orange), 2 (light blue), and 3 or more (dark blue) as determined by amplification of the hygromycin selectable marker. **A)** All individuals inoculated with *Pgt* race QTHJC showing most individuals with a susceptible score of > 8. **B)** Subset of the histogram for phenotypes less than 8 shown as an inset for clarity.

**Supplemental Figure 7.**
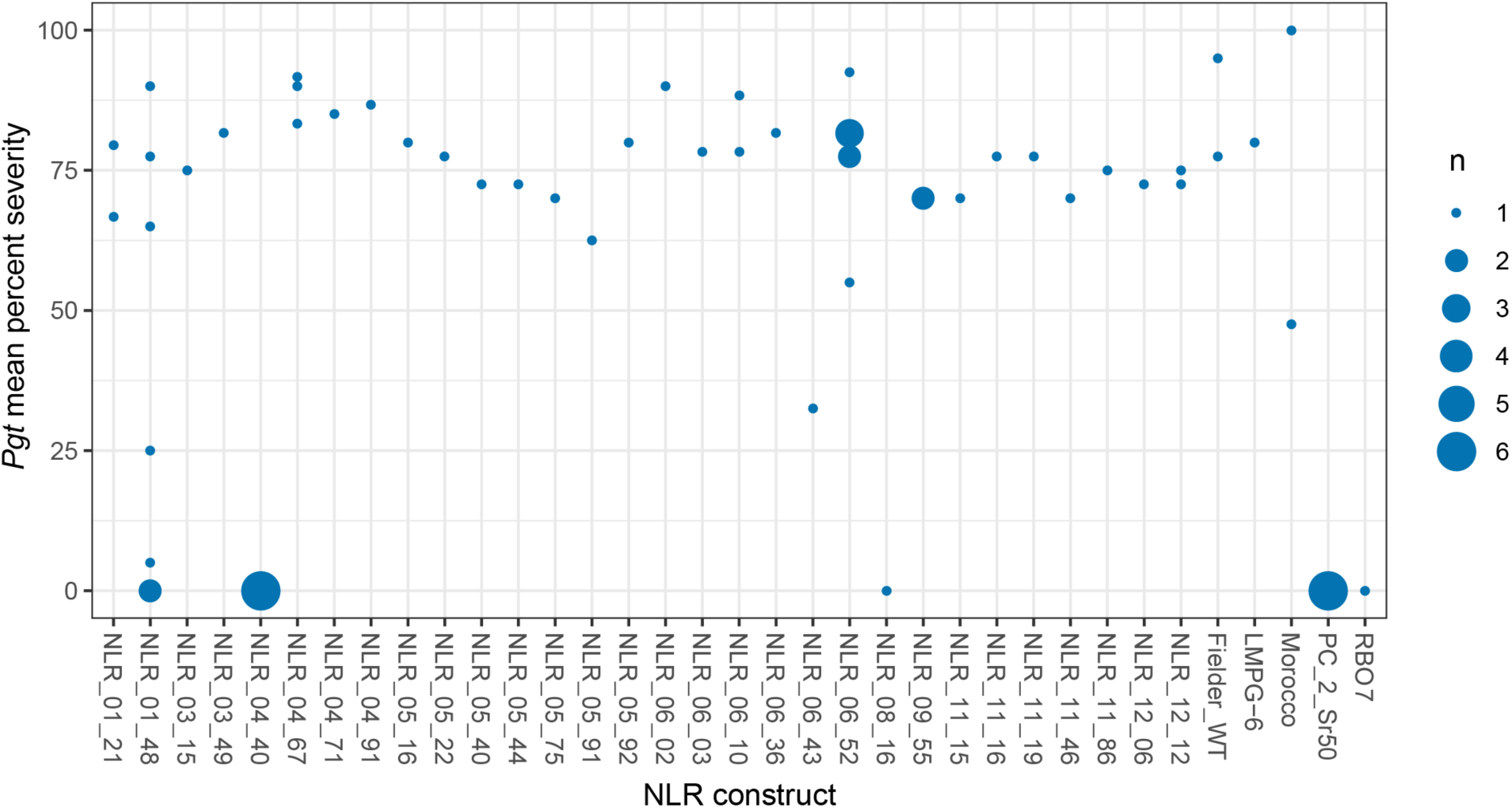
Field resistance of adult plants to *Puccinia graminis* f. sp. *tritici* race QTHJC. Phenotypic scores from individuals within T_2_ families from each construct inoculated with *Pgt* race QTHJC under field conditions during 2021 and 2022 (**Supplemental Data 5**). Circle size indicates number of individuals at each phenotypic score. Resistant controls included cultivar RBO7 and transgenic lines with *Sr50*. The cultivars Fielder, Morocco, LMPG-6 were included as susceptible controls. Phenotypes scored as percent severity recorded as the visual percentage (0-100%) of tissue covered by uredinia. The lowest phenotype was used from segregating lines due to assumed segregation of the transgene. The average of the replicates per transgenic line is shown. Individuals from T_2_ families of the constructs NLR_01_48, NLR_04_40, NLR_06_43, and NLR_08_16 displayed resistance with a lower percent severity phenotype as compared to the wild-type Fielder (Fielder WT).

**Supplemental Figure 8.**
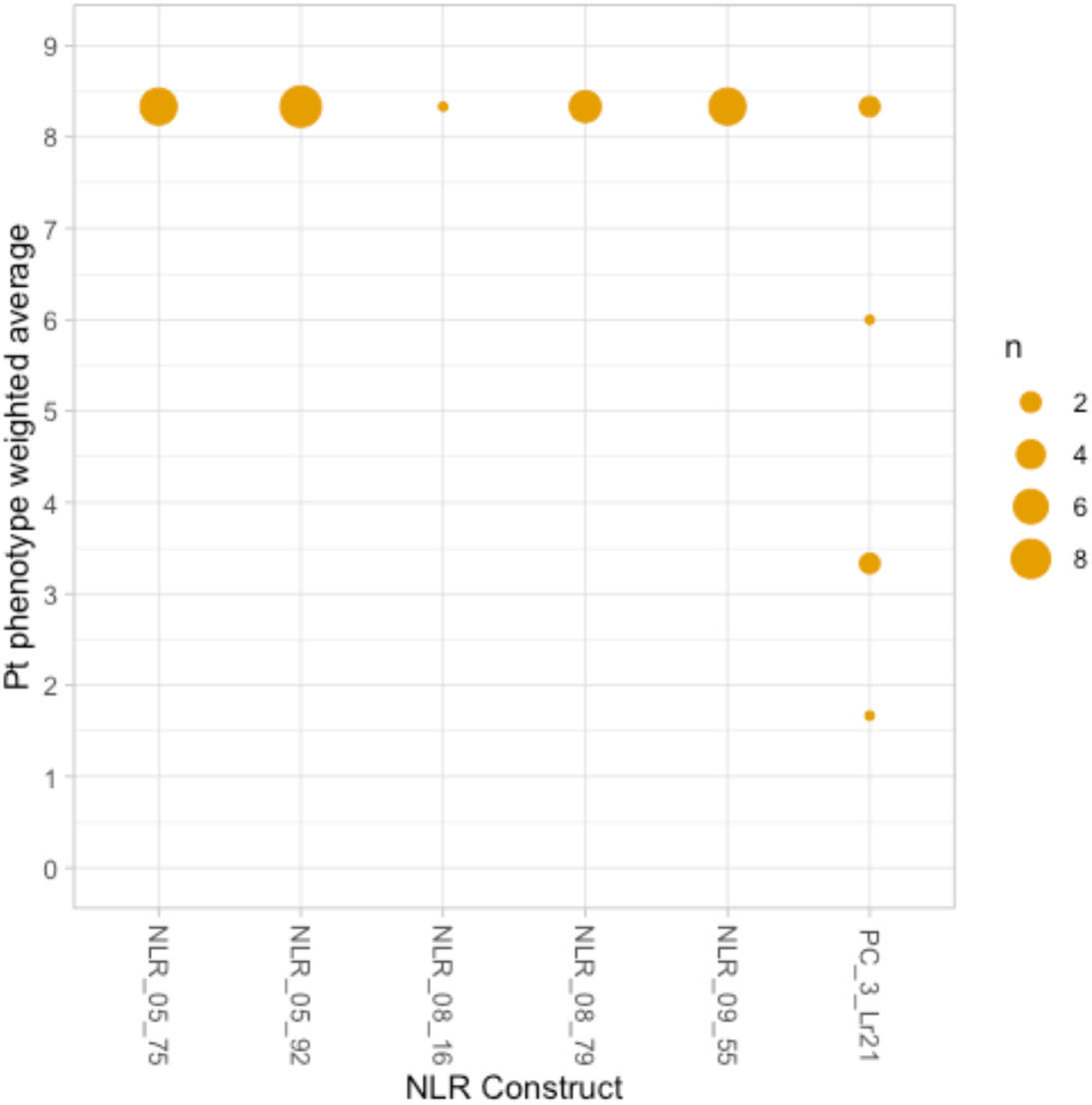
NLRs providing resistance to stem rust are susceptible to wheat leaf rust. Phenotypic scores from individuals within T_1_ families from each construct inoculated with *Puccinia triticina* race THBJ plotted on a weighted and transformed scale from completely resistant (0) to completely susceptible (9). Circle size indicates number of individuals having each phenotypic score.

**Supplemental Figure 9.**
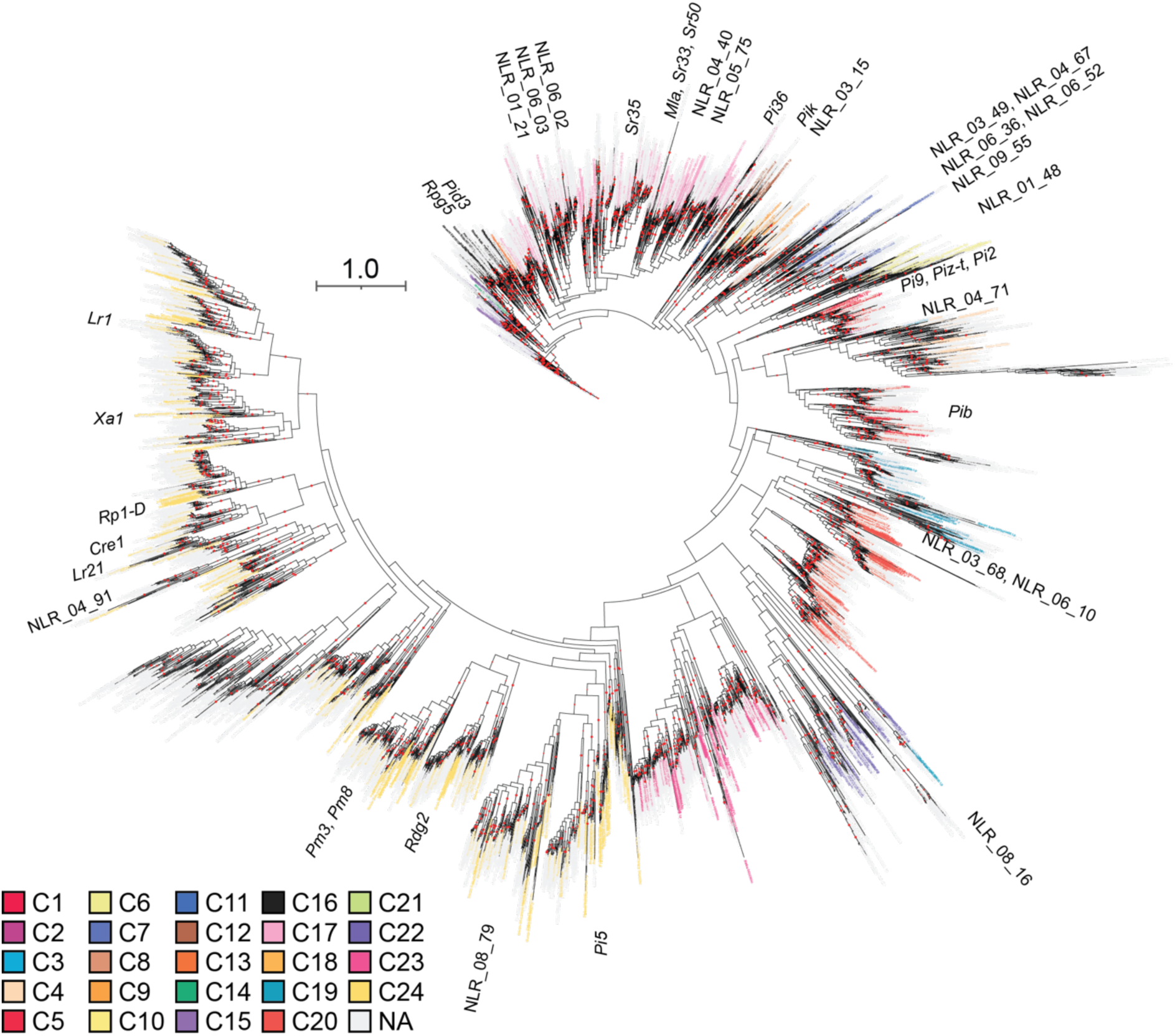
Phylogenetic tree of diverse grass species NLR repertoire and resistant NLRs. The maximum likelihood phylogenetic tree is based on the NB domain of NLRs from *Aegilops tauschii*, *Brachypodium distachyon*, *Hordeum vulgare* (barley), *Oryza sativa* (rice), *Setaria italica*, *Sorghum bicolor*, *Triticum aestivum* (wheat), and *Triticum urartu*, *Zea mays* (maize), previously cloned NLRs, and resistant NLRs in this work using the approach and clade classification of Bailey *et al*. (2018). NLR identifiers are labelled based on clade classification (key in bottom right). Bootstrap support of greater than 80% is indicated by orange dots on branches based on 1,000 bootstraps. The phylogenetic tree contains 4,990 non-redundant NB domains from NLRs.

**Supplemental Table 1.** The number of accessions sequenced and the number of NLRs cloned per species.

| Species | Acronym | Accessions | NLRs |
| --- | --- | --- | --- |
| *Achnatherum hymenoides* | *Ahy* | 3 | 20 |
| *Aegilops bicornis* | *Abi* | 6 | 98 |
| *Aegilops longissima* | *Alo* | 5 | 79 |
| *Aegilops searsii* | *Aes* | 3 | 53 |
| *Aegilops sharonensis* | *Ash* | 16 | 227 |
| *Agropyron cristatum* | *Agc* | 2 | 32 |
| *Avena abyssinica* | *Ava* | 3 | 51 |
| *Brachypodium distachyon* | *Bdi* | 2 | 25 |
| *Briza media* | *Brm* | 3 | 55 |
| *Cynosurus cristatus* | *Ccr* | 3 | 39 |
| *Echinaria capitata* | *Ecc* | 1 | 10 |
| *Holcus lanatus* | *Hla* | 3 | 48 |
| *Hordeum vulgare* | *Hvu* | 5 | 39 |
| *Koeleria macrantha* | *Kma* | 3 | 65 |
| *Lolium perenne* | *Lop* | 3 | 50 |
| *Melica ciliata* | *Mci* | 2 | 51 |
| *Phalaris coerulescens* | *Pco* | 3 | 11 |
| *Poa trivialis* | *Ptr* | 3 | 66 |

**Supplemental Table 2.**
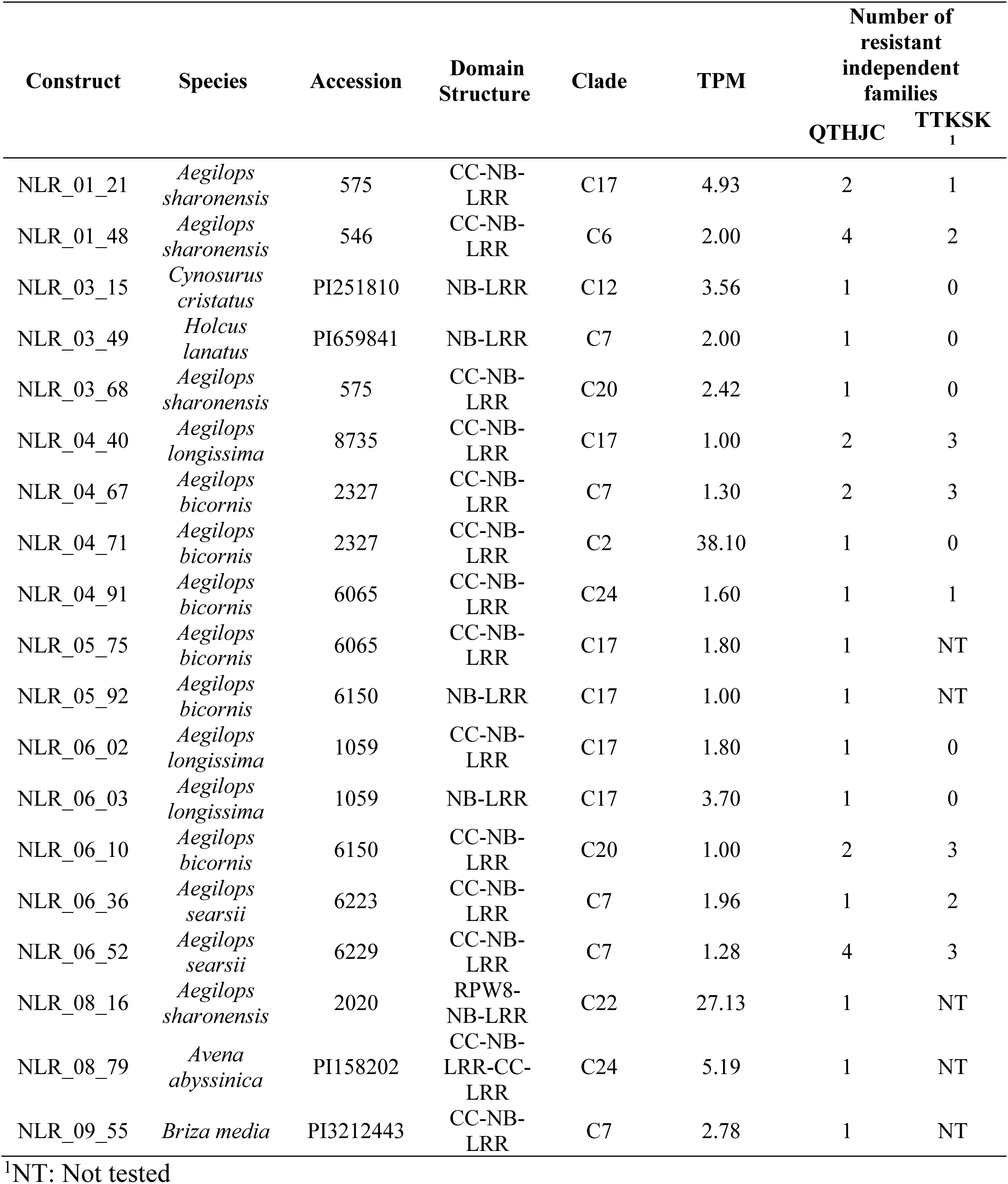
Resistant NLRs against the wheat stem rust (*Puccinia graminis* f. sp. *tritici; Pgt*) following screening of transgenic T_1_ families with *Pgt* races QTHJC and TTKSK.

